# Combining 3D single molecule localization strategies for reproducible bioimaging

**DOI:** 10.1101/385799

**Authors:** Clément Cabriel, Nicolas Bourg, Pierre Jouchet, Guillaume Dupuis, Christophe Leterrier, Aurélie Baron, Marie-Ange Badet-Denisot, Boris Vauzeilles, Emmanuel Fort, Sandrine Lévêque-Fort

## Abstract

We developed a 3D localization-based super-resolution technique providing a slowly varying localization precision over a 1 μm range with precisions down to 15 nm. The axial localization is performed through a combination of point spread function (PSF) shaping and supercritical angle fluorescence (SAF), which yields absolute axial information. Using a dual-view scheme, the axial detection is decoupled from the lateral detection and optimized independently to provide a weakly anisotropic 3D resolution over the imaging range. This method can be readily implemented on most homemade PSF shaping setups and provides drift-free, tilt-insensitive and achromatic results. Its insensitivity to these unavoidable experimental biases is especially adapted for multicolor 3D super-resolution microscopy, as we demonstrate by imaging cell cytoskeleton, living bacteria membranes and axon periodic submembrane scaffolds. We further illustrate the interest of the technique for biological multicolor imaging over a several-μm range by direct merging of multiple acquisitions at different depths.

---

Despite recent advances in localization-based super-resolution techniques, nanoscale 3D fluorescence imaging of biological samples remains a major challenge, mostly because of its lack of versatility. While photoactivated localization microscopy (PALM) and (direct) stochastic optical reconstruction microscopy ((d)STORM) can easily provide a lateral localization precision (i.e. the standard deviation of the position estimates) down to 5–10 nm [1, 2, 3, 4], a great deal of effort is being made to develop quantitative and reproducible 3D super-localization methods. The most widely used 3D Single Molecule Localization Microscopy (SMLM) technique is astigmatic imaging, which relies on the use of a cylindrical lens to apply an astigmatic aberration in the detection path to encode the axial information in the shape of the spots, achieving an axial localization precision (standard deviation) down to 20–25 nm [5]—though the precision sharply varies with the axial position: 300 nm away from the focus, the precision is typically around 60 nm (see **Supplementary Fig. 1a**). Other Point Spread Function (PSF) shaping methods are also available [6, 7, 8], but their implementations are not as inexpensive and straightforward. Still, all PSF shaping methods including astigmatic imaging suffer from several bias sources such as axial drifts, chromatic aberrations, field-varying geometrical aberrations and sample tilts. These sources of biases often degrade the resolution or hinder colocalization and experiment reproducibility. Axial measurements can also be performed thanks to intensity-based techniques like Supercritical Angle Fluorescence (SAF) [9, 10, 11, 12, 13, 14], which relies on the detection of the near-field emission of fluorophores coupled into propagative waves at the sample/glass coverslip interface due to the index mismatch. Combined with SMLM, this technique, called Direct Optical Nanoscopy with Axially Localized Detection (DONALD) or Supercritical Angle Localization Microscopy (SALM), yields absolute axial positions (i.e. independent of the focus position) in the first 500 nm beyond the coverslip with a precision down to 15 nm [15, 16]. The principle relies on the comparison between the SAF and the Undercritical Angle Fluorescence (UAF) components to extract the absolute axial position.

We set out to develop a microscopy technique that offers precise and unbiased results to enable reliable quantitative biological studies in the first micron beyond the coverslip. Starting from the efficient and straightforward astigmatic imaging, we propose to push back its previously mentioned limits thanks to a novel approach based on a dual-view setup (**Fig. 1a**) that combines two features. First, it decouples the lateral and axial detections to optimize the 3D localization precision, and second, it uses two different sources of axial information: a strong astigmatism-based PSF measurement is merged with a complementary SAF information that provides an absolute reference. This reference is crucial to render the axial detection insensitive to axial drifts and sample tilts as well as chromatic aberrations: unlike most other techniques that use fiducial markers [17] or structure correlation [5] to provide these corrections, here we intend to use the fluorophores themselves as absolute and bias-insensitive references. Besides, by applying a large astigmatic aberration on one fluorescence path only, this technique optimizes the axial precision for the collected photon number (**Supplementary Fig. 1b**) and maintains a slowly varying localization precision over the imaging depth (**Supplementary Fig. 1a**). Unlike most PSF shaping implementations found in the literature, which use moderate aberrations [5, 18, 19] to preserve the lateral resolution, the dual path detection allows one to fully benefit from the astigmatism capabilities. Indeed, as the lateral detection is mostly provided by the aberration-free path, the strong PSF shaping does not compromise the lateral detection. In order to merge the axial and lateral information sources, each is assigned a relative weight according to its localization precision (see **Fig. 1b** and **Methods**). Such a setup exhibits a major improvement in terms of both axial precision and precision curve flatness despite only half of the photons being used for the axial localization far from the coverslip compared to a standard single-view PSF measurement microscope. As a result, this technique, called Dual-view Astigmatic Imaging with SAF Yield (DAISY), exhibits a weakly anisotropic resolution over the whole capture range. We first performed the calibration of the astigmatism-based axial detection using 15 μm diameter latex microspheres coated with Alexa Fluor (AF) 647 as described in [20] in order to account for the influence of the optical aberrations on the PSFs and thus eliminate this axial bias source (see **Methods**). Then, to evaluate the localization precision of DAISY, we imaged dark red 40-nm diameter fluorescent beads located at various randomly distributed heights with a weak 637 nm excitation so that their emission level matched that of AF647 in typical dSTORM conditions, i.e. 2750 UAF photons and 2750–5100 EPI photons (dependeing on the depth) per bead per frame on average (**Fig. 1c**). As it takes advantage of the good performance of the SAF detection near the coverslip, DAISY exhibits a resolution that slowly varies with depth: the lateral and axial precisions reach values as low as 8 nm and 12 nm respectively (standard deviations), and they both remain inferior to 20 nm in the first 600 nm. Such precision is sufficient to resolve the hollowness of immunolabeled microtubules, as displayed in **Supplementary Fig. 2**. This feature is rather uncommon with astigmatic imaging implementations, which typically provide at best 20–25 nm axial precision [5] and only in a limited axial range of approximately 300 nm according to Cramér-Rao Lower Bound (CRLB) calculations (**Supplementary Fig. 1a**, **Supplementary Fig. 3**)—only the dual-objective implementation achieves better precisions, at the cost of a much increased complexity [21]. It is worth noticing that the experimental precisions are slightly superior to the CRLB, which represent a theoretical ideal. This discrepancy is most likely due to optical aberrations, which are not taken into account by the CRLB, and to the use of centroid detection (see **Methods**), which is not expected to reach the lower limit.

**Figure 1:**
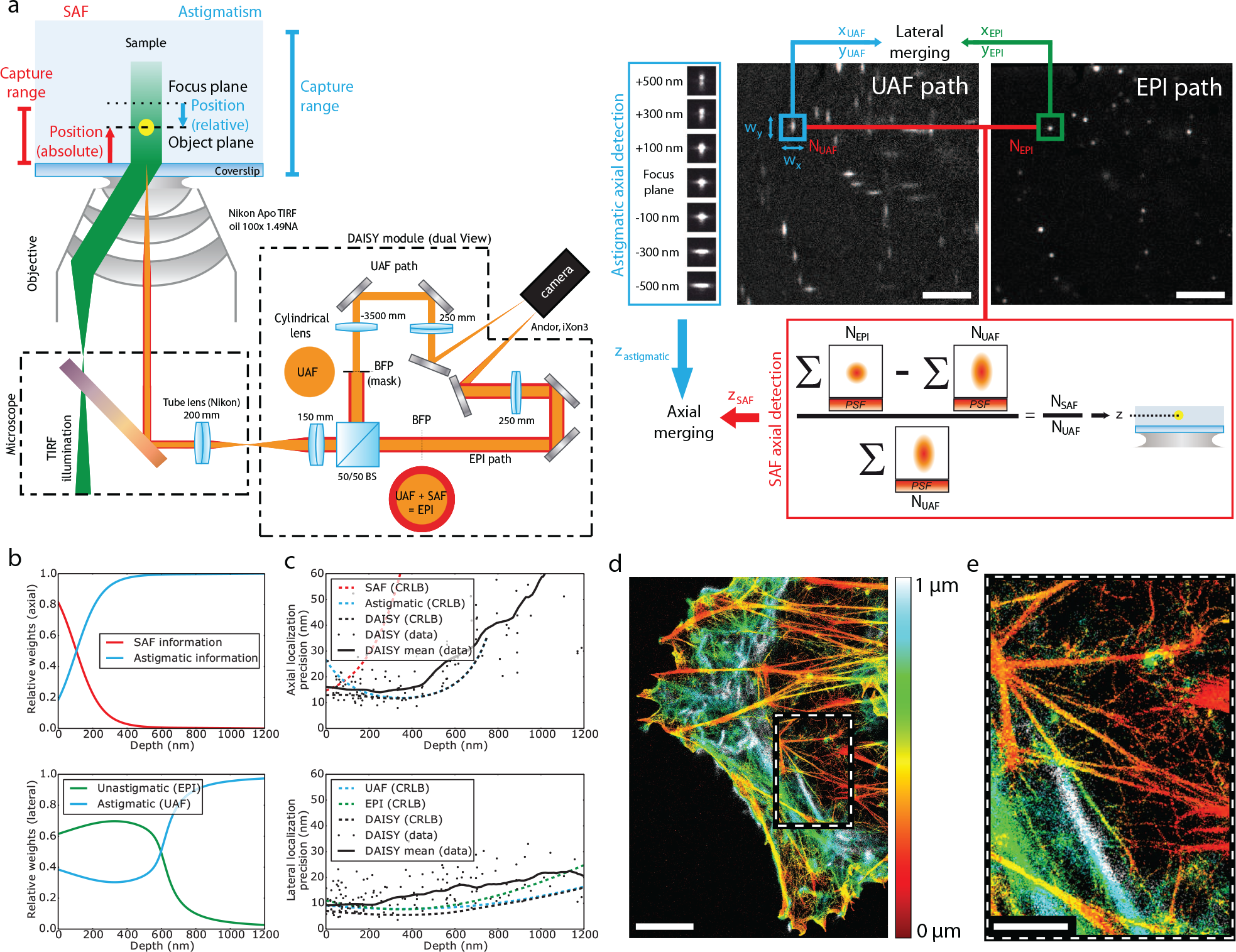
Description of the principle of DAISY and characterization of the precision. **(a)** Schematic of the setup. The DAISY module is placed between the microscope and the camera. After the beam splitter cube (BS), the Undercritical Angle Fluorescence (UAF) path contains a cylindrical lens, as well as a physical mask in a relay plane of the back focal plane of the objective to block the SAF photons. These two elements are not present in the epifluorescence (EPI) detection path, which comprises both the UAF and SAF components. The images are formed on the two halves of the same camera. UAF and EPI frames recorded by the camera on a given field (COS-7 cells, *α*-tubulin immunolabeling, Alexa Fluor 647) are also displayed (top right corner). For each PSF, the *x* and *y* widths are measured to obtain the astigmatic axial information, and the numbers of UAF and EPI photons are used to retrieve the SAF axial information. Finally, the axial astigmatic and SAF positions are merged together. Similarly, lateral positions are obtained by merging the lateral positions from the UAF and EPI paths. **(b)** Relative weights of the SAF and astigmatic axial detections (top) and of the UAF and EPI lateral positions (bottom) used to merge the positions in DAISY (see **Methods**, **Position merging** section for the exact formulas). **(c)** Axial (top) and lateral (bottom) precisions of DAISY. The experimental data was taken on dark red 40-nm fluorescent beads distributed at various depths, each emitting a number of photons similar to Alexa Fluor 647. 500 frames were acquired and the precisions were evaluated from the dispersion of the results for each bead. The CRLB contributions of each detection modality are also displayed, as well as the CRLB of DAISY for typical experimental conditions. **(d)** 3D (color-coded depth) DAISY image of actin (COS-7 cell, AF647-phalloidin labeling). **(e)** Zoom on the boxed region displayed in **(d)**. Scale bars: 5 μm **(a)** and **(d)**, 2 μm **(e)**.

Our technique thus provides precise 3D super resolution images (**Fig. 1d–e**); still, at this precision level, any experimental uncertainty or bias can have devastating effects on the quality of the obtained data. The first source of error that has to be dealt with is the drifts that typically come from a poor mechanical stability of the stage or from thermal drifts. Lateral drifts are well known and can often be easily corrected directly from the localized data using cross-correlation algorithms [22]. However, accounting for the axial drifts can be much more demanding since 3D cross-correlation algorithms require long calculation times unless they sacrifice precision. Tracking fiducial markers is also possible, but since it requires a specific sample preparation and is sensitive to photobleaching (unless a dedicated detection channel at a different wavelength is used [17]), it is not very practical. It is worth noticing that most commercially available locking systems typically stabilize the focus position at ±30 nm at best (**Supplementary Fig. 4**), which is hardly sufficient for high resolution imaging. As positions are measured relative to the focus plane with PSF shape measurement methods, axial drifts induce large losses of resolution. On the contrary, SAF detection yields absolute results; thus it is not sensitive to drifts. We use this feature to provide a reliable drift correction algorithm: for each localization, the axial position detected with the SAF and the astigmatic modalities are crosscorrelated, which allows us to monitor the focus drift and to consequently correct the astigmatism results with an accuracy typically below 6 nm (see **Methods**). To highlight the importance of this correction, we plotted the *x*-*z* and *y*-*z* profiles of a microtubule labeled with AF647 as a function of time with both an astigmatism-based detection and DAISY (**Fig. 2a**): unlike the DAISY profiles, the astigmatism profiles exhibit a clear temporal shift, which results in a dramatic apparent broadening of the filament.

**Figure 2:**
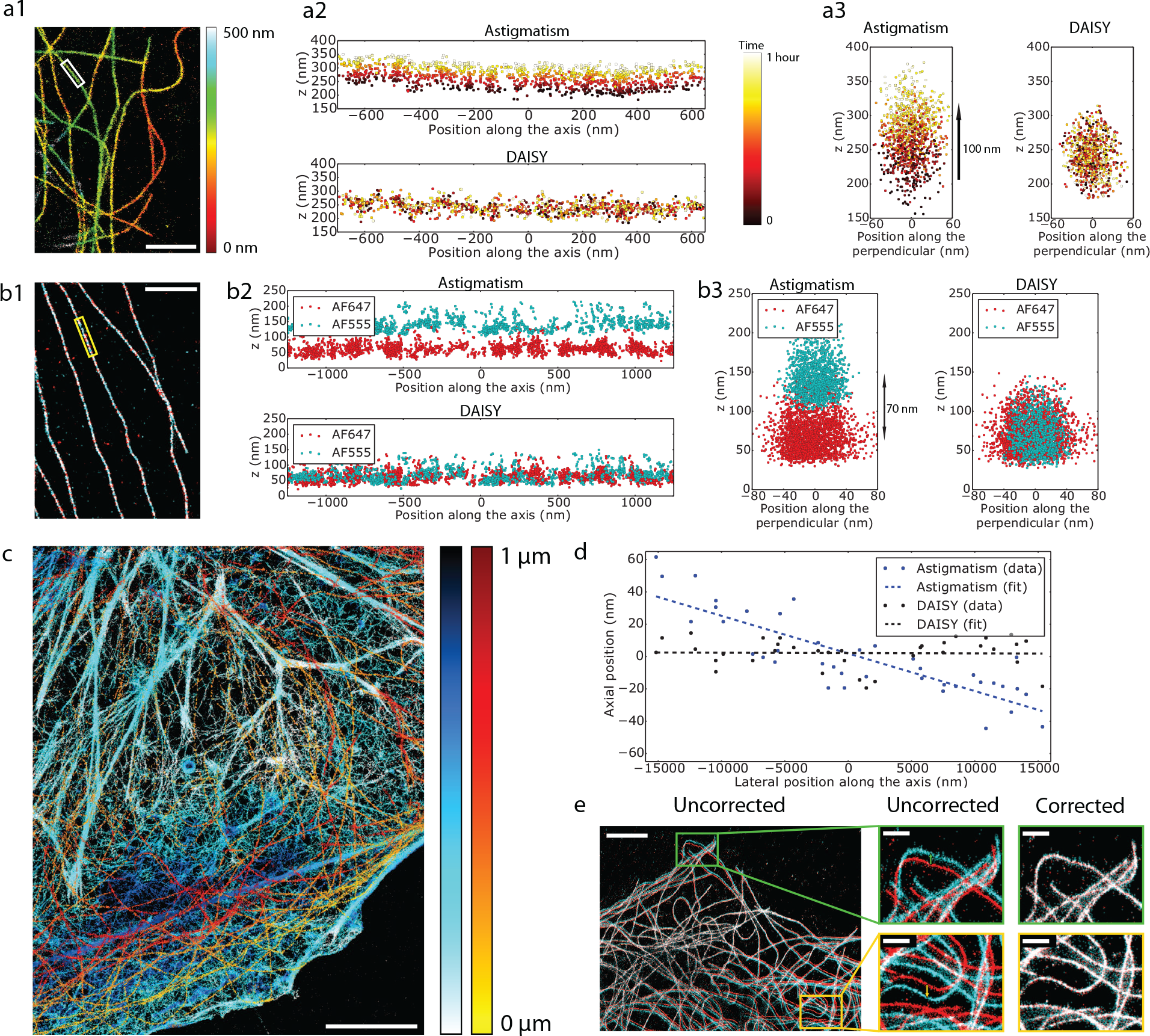
Characterization of the performance of DAISY. **(a)** Illustration of the effect of axial drifts. **(a1)** depth map of microtubules (COS-7 cells, *α*-tubulin labeled with AF647). The *x*-*z* **(a2)** and *y*-*z* **(a3)** profiles of the boxed microtubule are plotted for both standard astigmatic imaging and DAISY. The time is color-coded over one hour to highlight the effect of the temporal drift. **(b)** Effect of the chromatic aberration. **(b1)** 2D localization image of microtubules (COS-7 cells, *α*-tubulin labeled with AF555 and *β*-tubulin labeled with AF647) sequentially imaged in two different colors (red: AF647, cyan: AF555). The *x*-*z* **(b2)** and *y*-*z* **(b3)** profiles of the boxed microtubule are plotted for both standard astigmatic imaging and DAISY. **(c)** Dual-color depth map of actin (cyan-blue) and tubulin (yellow-red) in COS-7 cells (actin labeled with AF647-phalloidin and *α*-tubulin labeled with a 560-nm excitable DNA-PAINT imager). **(d)** Influence of the sample tilt on the axial detection. The same field of 20-nm dark red fluorescent beads deposited on a coverslip was imaged with both standard astigmatic imaging and DAISY and the results were averaged over 500 frames to suppress the influence of the localization precision. The detected depth profile is plotted along the tilt axis. **(e)** Illustration of the image lateral deformation induced by the astigmatism. For the same acquisition (COS-7 cells, *α*-tubulin labeled with AF647), 2D images were reconstructed from the lateral positions measured on both the astigmatic UAF (in cyan) and the unastigmatic EPI (in red) detection paths of our setup, before the deformation correction (left) and after (right). The whole field and zooms on the boxed regions are both displayed. Scale bars: 2 μm **(a1)** and **(b1)**, 5 μm **(c)** and **(e)** left, 1 μm **(e)** right insets.

In the framework of quantitative biological studies, the axial detection can furthermore be hampered by the axial chromatic aberration due to dispersion by the lenses, including the objective lens. If uncorrected, such a chromatic shift induces a bias in the results of multicolor sequential acquisitions, thus hindering colocalization. However, as DAISY provides absolute axial information thanks to the SAF measurement, it is not sensitive to this chromatic aberration. We performed a two-color sequential acquisition on microtubules labeled with AF647 and AF555 (**Fig. 2b**). It illustrates the chromatic dependence inherent in standard PSF shaping detection (which exhibits chromatic shifts as large as 70 nm) and the insensitivity of DAISY to this effect (residual chromatic shift inferior to 5 nm). Because of the chromatic shift, the uncorrected astigmatism results appear somewhat inconsistent, whereas the colocalization is much more obvious with DAISY. Consequently, unbiased dual-color 3D images of biological samples can be obtained thanks to sequential acquisitions: we illustrate this on a sample with the actin and the tubulin labeled with AF647 and a 560-nm-excitable DNA-PAINT fluorophore respectively (**Fig. 2c**).

It is well known that axial biases in PSF shaping measurements can further stem from tilts of the stage or sample holder, as well as from field-dependent geometrical optical aberrations. These issues were thoroughly studied by Diezmann *et al.*, who reported discrepancies higher than 100 nm over one field of view [23]. Although assessing tilts on biological samples is difficult with PSF measurement methods, DAISY makes this measurement straightforward since the absolute reference provided by the SAF detection can be used to measure the values of the astigmatic axial positions detected for molecules at the coverslip as a function of their lateral positions and then correct the tilt. We performed DAISY acquisitions on 20-nm diameter fluorescent beads at the coverslip and displayed the *z* values obtained with both an astigmatism-based detection and DAISY. While the former exhibits a clear tilt ranging from −30 nm to +30 nm over a 30 μm wide field, the latter is insensitive to the tilt, with less than 2 nm axial discrepancy between the two sides of the field (**Fig. 2d**).

Aside from tilt effects, field-dependent aberrations also induce PSF shape deformations, leading to axial biases. Although we do not actually perform corrections, DAISY is less sensitive to that effect compared to standard astigmatism imaging: on the one hand, the SAF detection relies on intensity measurement, and on the other hand, as DAISY uses a high astigmatism, i.e. strongly aberrated PSFs, it exhibits little sensitivity to remaining field aberrations. To illustrate this phenomenon, we compared tilt-corrected axial positions obtained with 20-nm diameter fluorescent beads deposited on a coverslip between a standard weaker astigmatic detection (350 nm between the two focal lines, close to the values commonly found in the literature) and DAISY. We got rid of the dispersion due to the localization precision by averaging the results over time for each bead and we plotted the corresponding detected depth histograms over one 25-μm wide field of view (**Supplementary Fig. 5**). The widths of the distributions evidence a much lower impact on the DAISY detection (standard deviation equal to 21 nm) than on the standard astigmatic detection (standard deviation equal to 45 nm). In other words, the strong astigmatism is less sensitive to aberrations than a conventional astigmatism, and the biases are even further mitigated by the coupling with the SAF detection, which relies on photon counting, and is thus weakly sensitive to PSF shapes.

To illustrate the accuracy of the axial correction of the astigmatism data using the SAF measurement, we performed measurements on 40-nm fluorescent beads, both at the coverslip and distributed in the volume (**Supplementary Fig. 6**). In both cases, the axial correction algorithm seems very accurate (1 nm average discrepancy at the coverslip, and 3 nm in the volume, which is well below the localization precision). As the dispersion of the values increases for beads in the volume, this can be attributed to either the decay of the SAF signal in the volume, which causes the SAF localization precision to become non-negligible, or the influence of the previously mentioned field-dependent aberrations, which induce biases in the astigmatic positions according to the position in the field. This effect is present in conventional single-view PSF shaping imaging too, but it is difficult to detect unless a specifically designed calibration sample is used. The dispersion due to field-dependent aberrations could be mitigated by using a spatially resolved PSF calibration, as in [23].

Lastly, the optical aberrations applied in PSF shaping-based setups not only deform the PSFs, but they may also distort the field itself laterally. For instance, when astigmatism is used, the system has two different focal lengths in *x* and *y*, which implies that the magnification is different in *x* and *y*. While this effect is of the order of a few percent, it definitely biases the results whenever it is necessary to measure lateral distances precisely unless this magnification discrepancy is duly calibrated. With DAISY, evaluating this image distortion is straightforward thanks to the non-astigmatic detection path: a cross-correlation performed between the astigmatic (UAF) and the unaberrated (EPI) 2D SMLM images gives the optimal affine transformation to be applied to the astigmatic image—this combination of translation, rotation and magnification directly provides the magnification difference between the *x* and *y* axes, which accounts for 3.5% approximately in our case (**Fig. 2e**). By applying the optimal affine transformation, the deformation is then corrected: the final lateral discrepancy between the two images was found to be below 6 nm over the whole 25 nm-wide field in **Fig. 2e** (see **Supplementary Fig. 7** for a more detailed measurement of the registration error). It should be noticed, however, that a solution to avoid such a deformation would be to place the cylindrical lens in the Fourier plane, although most reported PSF shaping setups do not use this optical configuration. Also, more complex PSF shapes might induce complex field distortions—potentially making the correction more difficult.

To evidence the performance of DAISY for unbiased, reproducible and quantitative experiments, we used it to image biological samples. We illustrate the performances in terms of resolution by performing acquisitions on living *E. coli* bacteria adhered to a coverslip. The envelope of bacteria was labeled with both AF647 and AF555 using a click chemistry process (see **Methods**) [24, 25]. Since the lipopolysaccharide (LPS) layer is thin in Gram-negative bacteria, this is a good sample to observe the influence of the localization precision. We present in **Fig. 3a–b** 2D and 3D images of a region of interest and in **Fig. 3c** an *x*-*z* slice along the line displayed in **Fig. 3a**. The measured diameter of the bacterium is around 1 μm but still it does not exhibit a strong loss of resolution at its edges. To evidence this, we also plotted the lateral and axial histograms in the boxed regions (**Fig. 3c**). The axial standard deviations were found to be respectively around 30 nm and 45 nm at the bottom and at the top of the cell, while lateral standard deviations were around 27 nm in both colors. Taking into account the size of the LPS layer (<10 nm), of the label—i.e. the DBCO-sulfo-biotin and streaptavidin-AF construction—(10 nm) and the effect of the curvature of the bacterium over the width of the area used for the analysis (10 nm), these values are consistent with the localization precision curves plotted in **Fig. 1c**. As a comparison, the results obtained on the same sample with uncorrected astigmatism and with DONALD are provided in **Supplementary Fig. 8**. Like DAISY, DONALD features an absolute detection, unsensitive to both chromatic aberration and axial drift. However, the axial precision of DONALD deteriorates sharply with the depth due to the decay of the SAF signal; thus the top half of the sample (beyond 500 nm) is hardly visible, whereas DAISY clearly permits imaging up to 1 μm. Uncorrected astigmatism has the same capture range as DAISY, but since it lacks the absolute information, it exhibits an axial shift between the two colors as well as a broadening of the histogram widths due to the axial drift.

**Figure 3:**
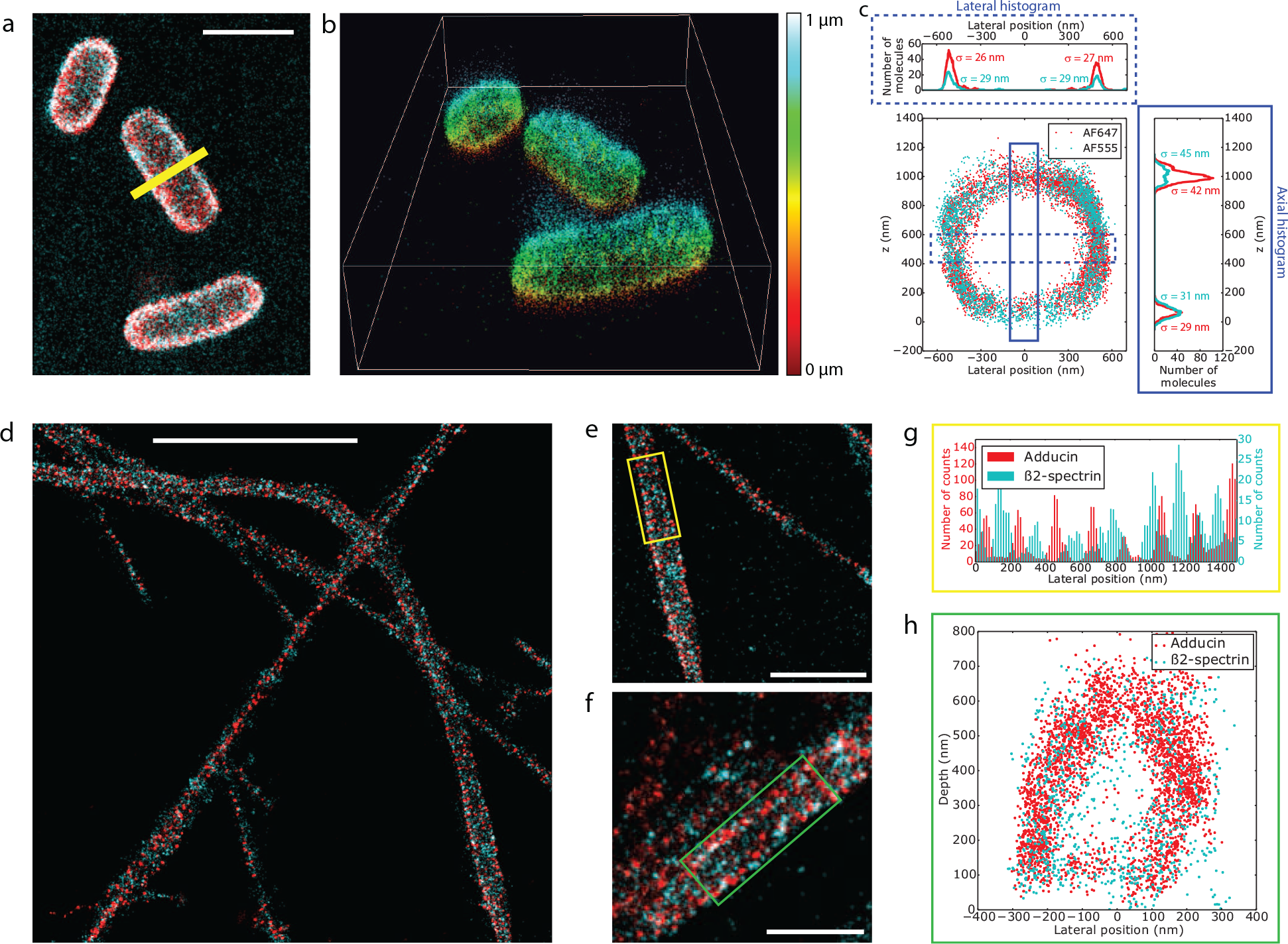
DAISY results obtained from biological samples. **(a)** 2D SMLM image of living *E. coli* bacteria labeled with both AF647 (red) and AF555 (cyan) at the membrane. **(b)** 3D view of the field displayed in **(a)**. The depth is color-coded (one single colormap is used for both AF647 and AF555). **(c)** *x*-*z* slice along the line displayed in **(a)** and axial and lateral profiles in the boxed regions. The *σ* values stand for the standard deviations of the distributions. **(d–f)** 2D dual-color images of rat hippocampal neurons where the adducin and the *β*2-spectrin were labeled with AF647 and AF555 respectively. **(g)** Lateral profile along the axis of the yellow box displayed in **(e)**. **(h)** *x*-*z* slice along the green box displayed in **(f)**. Scale bars: 2 μm **(a)** and **(e)**, 5 μm **(d)**, 1 μm **(f)**.

We then used DAISY to visualize the periodic submembrane scaffold present along the axon of cultured neurons [26]. We imaged the 3D organization of two proteins within this scaffold: adducin (labeled with AF647) that associates with the periodic actin rings, and *β*2-spectrin (labeled with AF555) that connect the actin rings (**Fig. 3d–f**). The lateral resolution allowed us to easily resolve the alternating patterns of adducin rings and *β*2-spectrin epitopes and their 190 nm periodicity (**Fig. 3g**) [27]. Thanks to the axial resolution of DAISY, we were also able to resolve the submembrane localization of both proteins across the whole diameter of the axon at 600 nm depth (**Fig. 3h**).

Taking advantage of the features of DAISY for unbiased sequential imaging, we propose an implementation allowing single-color and multicolor imaging at wider depth ranges by stacking the results of multiple acquisitions on the same field at different heights. Although PSF measurement methods also allow this type of acquisitions, DAISY is especially suited in this case thanks to its previously described intrinsic bias correction features. Since the SAF signal quickly decays with the depth in the first 500 nm above the coverslip, the absolute reference is accessible only in the first stack. Still, as it provides unbiased results, the top of this first stack serves as an absolute reference for the next stack, which is matched to the previous using an axial position cross-correlation algorithm. In other words, the first 1 μm unbiased slice is interlaced with the following one, which contains the positions between 600 nm and 1.6 μm (as described in the schematic in **Fig. 4a**). The absolute reference is thus transferred from the first slice onto the second, which becomes insensitive to axial detection biases. Similarly, the third slice, containing positions from 1.2 μm to 2.2 μm is intertwined with the second by position cross-correlation and thus it also benefits from the absolute reference and the bias insensitivity that it brings. Several slices can be recorded and merged together to obtain an extended depth image—still, this is limited by photobleaching (although this can be mitigated by using (DNA-)PAINT labeling), as well as aberrations inherent in depth imaging, which cause the axial and lateral precisions to deteriorate away from the coverslip. Moreover, registration errors are likely to add as the number of slices increases, so using fiducial markers might be necessary to merge more slices. We illustrate the method with a single-color acquisition series (COS-7 cells, *α*- and *β*-tubulin labeled with AF647) in **Fig. 4b–d**: the stack of the three slices (**Fig. 4e**) obviously shows information in deep regions (beyond 1 μm) that would not be accessible with a single acquisition. We then imaged a dual-label tubulin-clathrin sample (COS-7 cells, light chain and heavy chain clathrin labeled with AF647, *α*- and *β*-tubulin labeled with 560-nm-excitable DNA-PAINT imager) in three sequential acquisitions while shifting the focus by 600 nm between each of them to obtain a 3D dual-color 2 μm imaging range set of data (**Fig. 4f**). Aside from the fact that no axial mismatch between the subsequent acquisitions is observed, the localization precision remains satisfactory after 1.5 μm as it is limited only by the effect of the spherical aberration and sample-induced aberrations. To evidence this, we measured the dispersion of the localizations on two clathrin spheres located close to the ventral membrane (200 nm depth) and the dorsal membrane (1500 nm depth) respectively (**Fig. 4g–h**, **Supplementary Fig. 9**). The lateral and axial standard deviations were found to be 16 nm in *xy* and 17 nm in *z* at 200 nm depth, and 20 nm in *xy* and 27 nm in *z* at 1500 nm depth—as expected, the axial precision is more affected by the effect of the aberrations in the volume than the lateral precision.

**Figure 4:**
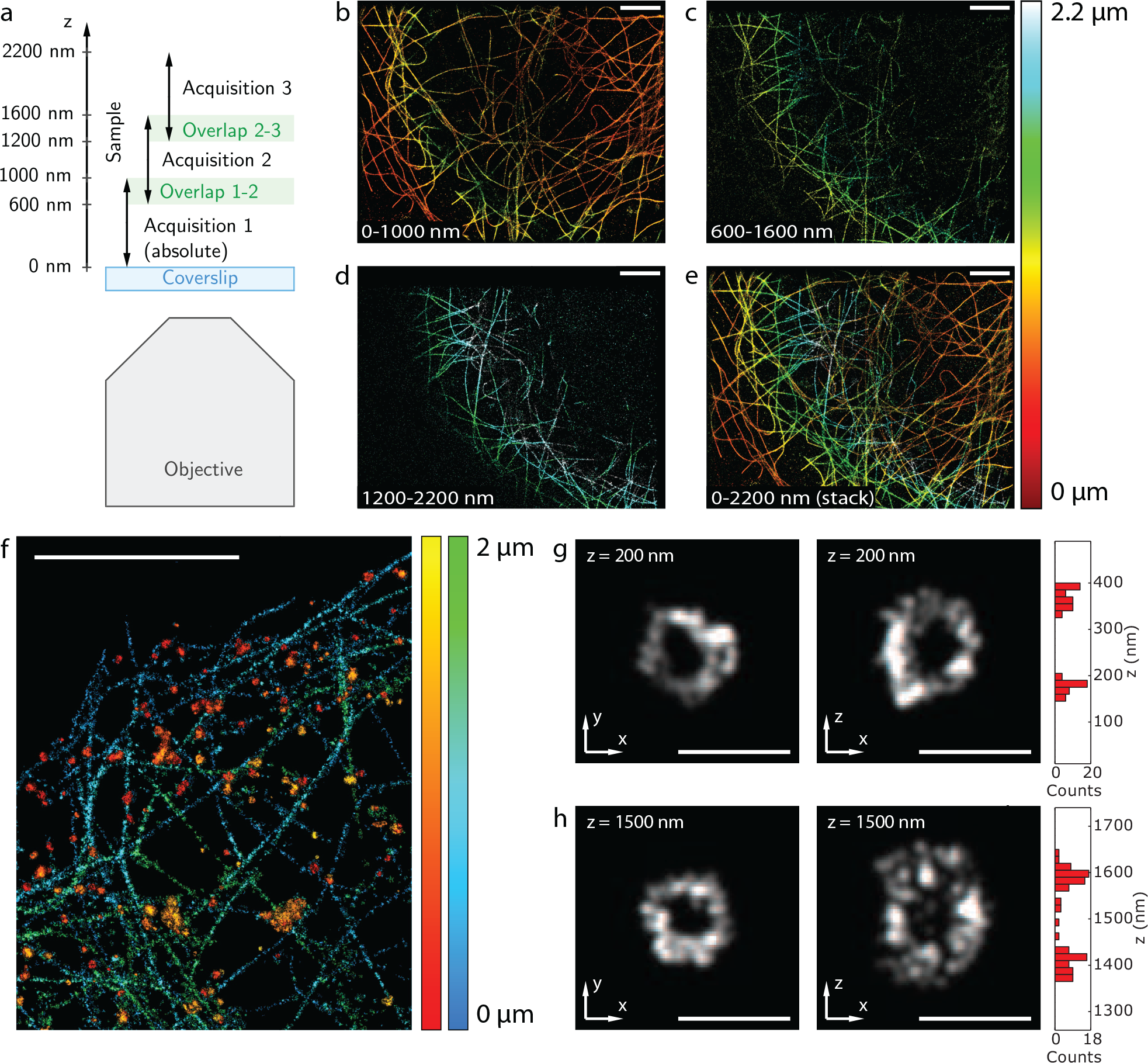
Extended depth imaging principle and results. **(a)** Description of the acquisition protocol: several sequential acquisitions are performed at different focus positions with a sufficient overlap between them to enable the stitching of the different slices (the focus is typically shifted by 600 nm between successive acquisitions, while the capture range is around 1 μm for each acquisition). **(b–d)** 3D images reconstructed from single-color tubulin acquisitions performed at three different focus positions (COS-7 cells, *α*- and *β*-tubulin labeled with AF647). **(e)** Final 3D image obtained by stitching the three consecutive acquisitions. The total range is around 2.2 μm. **(f)** 3D extended range dual-color image of clathrin (red-yellow) and tubulin (blue-green) obtained from three sequential acquisitions (for each color) at different heights (COS-7 cells, heavy chain and light chain clathrin labeled with AF647, *α*- and *β*-tubulin labeled with a 560-nm excitable DNA-PAINT imager). **(g–h)** *x*-*y* and *x*-*z* slices of two clathrin spheres taken from **(f)** at two different depths (200 nm and 1500 nm). The axial histograms of the *x*-*z* images are displayed on the right. Scale bars: 5 μm **(b–f)**, 250 nm **(g–h)**.

Thanks to the decoupling of the axial and lateral detections and to the combination of two axial SMLM techniques yielding complementary information, we could achieve reliable and unbiased imaging that enables quantitative studies on biological samples. DAISY offers a slowly varying, weakly anisotropic resolution over the whole micron-wide capture range, with a localization precision down to 15 nm. Thanks to both the SAF and the astigmatic detections, DAISY provides absolute axial results that prove to be insensitive to axial drifts and sample tilts as well as chromatic aberration. These features make it especially suited for biological samples imaging near the coverslip, which finds applications in the framework of cell adhesion, motility processes, bacteria imaging or neuronal axons and dendrites studies. Moreover, stacking acquisitions performed at different heights also enables reproducible and reliable studies at more important depths, up to a few μm. Finally, as the implementation of the dual-view detection scheme we use is straightforward, it would also benefit any PSF measurement method other than astigmatism, such as dual helix PSF [6], self-bending PSF [7], saddle-point PSF [8] and tetrapod [28], which offer better performances in terms of localization precision and capture range.

## Acknowledgements

We thank Ultivue for consumable gifts and Abbelight for software and buffers gifts.

We acknowledge the contribution of the Centre de Photonique BioMédicale to cell culture and labeling. We also acknowledge the help of Marion Bardou with cell culture. We thank Rym Boudjemaa for her contribution to the bacteria labeling project. Finally, we thank Caroline Schou and Yann Kergutuil for their help regarding software analysis.

This work was supported by the AXA research fund, the ANR (LABEX WIFI, ANR-10-LABX-24), the DIM CNANO Île-de-France, the IRS Bioprobe, the Mission interdisciplinarité of the CNRS and LaserLab-Europe EUH2020654148.

## Author contributions

C.C., N.B., P.J., G.D., E.F. and S.L.-F. conceived the project. C.C. designed the optical setup and performed the acquisitions. C.C. and N.B. carried out simulations and data analysis. P.J. and C.C. performed the CRLB calculations. N.B. developed the (d)STORM buffer. N.B., C.C. and P.J. optimized the immunofluorescence protocol. P.J. and C.C. prepared the COS-7 cells samples, C.L. prepared the neuron samples, A.B., M.-A. B.-D. and B.V. prepared the bacteria samples. All authors contributed to writing the manuscript.

## Competing financial interests

N.B., E.F. and S.L.F. are shareholders in Abbelight.

## Methods

### Optical setup

A schematic of the optical setup used is presented in **Fig. 1a**. We used a Nikon Eclipse Ti inverted microscope with a Nikon Perfect Focus System. The excitation was performed thanks to five different lasers: 637 nm (Obis 637LX, 140 mW, Coherent), 561 nm (Genesis MX 561 STM, 500 mW), 532 nm (Verdi G5, 5 W, Coherent), 488 nm (Genesis MX 488 STM, 500 mW, Coherent) and 405 nm (Obis 405LX, 100 mW, Coherent). The corresponding 390/482/532/640 or 390/482/561/640 multiband filters (LF405/488/532/635-A-000 and LF405/488/561/635-A-000, Semrock) were used. The fluorescence was collected through a Nikon APO TIRF x100 1.49 NA oil immersion objective lens, sent in the DAISY module and recorded on two halves of a 512×512-pixel EMCCD camera (iXon3, Andor). The camera was placed at the focal plane of the module of magnification 1.67 and the optical pixel size was approximately 100 nm. Finally, the imaging paths were calibrated in intensity to compensate the non-ideality of the 50-50 beam splitter as well as the reflection on the cylindrical lens surface (this measurement was performed for each fluorescence wavelength). The object focal plane of the EPI path was typically at the coverslip (*z* = 0 nm) and the UAF path had two focal lines, at *z* = 0 nm and *z* = 800 nm for the *y* and *x* axes respectively.

### Sample preparation

COS-7 cells were grown in DMEM with 10% FBS, 1% L-glutamin and 1% penicillin/streptomycin (Life Technologies) at 37°C and 5% CO_2_ in a cell culture incubator. Several days later, they could be plated at low confluency on cleaned round 25 mm diameter high resolution #1.5 glass coverslips (Marienfield, VWR). After 24 hours, the cells were washed three times with PHEM solution (60 mM PIPES, 25 mM HEPES, 5 mM EGTA and 2 mM Mg acetate adjusted to pH 6.9 with 1 M KOH) and fixed for 12 min in 4% PFA, 0.2% glutaraldehyde and 0.5% Triton; they were then washed 3 times in PBS (Invitrogen, 003000). Up to this fixation step, all chemical reagents were prewarmed at 37°C. The cells were post-fixed for 10 min with PBS + 0.1% Triton X-100, reduced twice for 10 min with NaBH_4_, and washed in PBS three times before being blocked for 15 min in PBS + 1% BSA.

The labeling step varied according to the required sample: in the case of actin labeling, the cells were incubated for 20 minutes with 3.3 nM phalloidin-AF647 (Thermo Fisher, A22287) in the (d)STORM imaging buffer (Abbelight) before starting the acquisition—without removing the (d)STORM buffer containing the phalloidin-AF647. On the contrary, immunolabeling of tubulin and clathrin required more preparation steps.

For AF647 *α*-tubulin, the cells were incubated for 1 hour at 37°C with 1:300 mouse anti-*α*-tubulin antibody (Sigma Aldrich, T6199) in PBS + 1% BSA. This was followed by three washing steps in PBS + 1% BSA, incubation for 45 min at 37°C with 1:300 goat anti-mouse AF647 antibody (Life Technologies, A21237) diluted in PBS 1% BSA and three more washes in PBS.

For AF647 *β*-tubulin and AF555 *α*-tubulin, the cells were incubated for 1 hour at 37°C with 1:300 rabbit anti-*β*-tubulin antibody (Sigma Aldrich, T5293) in PBS + 1% BSA. This was followed by three washing steps in PBS + 1% BSA, incubation for 45 min at 37°C with 1:300 goat anti-rabbit AF555 antibody (Life Technologies, A21430) diluted in PBS + 1% BSA and three more washes in PBS + 1% BSA. Then they were incubated again for 1 hour at 37°C with 1:300 mouse anti-*α*-tubulin antibody (Sigma Aldrich, T6199) in PBS + 1% BSA, washed three times, incubated for 45 min at 37°C with 1:300 goat anti-mouse AF647 antibody (Life Technologies, A21237) diluted in PBS + 1% BSA and washed three more washes in PBS.

For AF647 *α*- and *β*-tubulin, the cells were incubated for 1 hour at room temperature with 1:300 mouse *β*-tubulin antibody (Sigma Aldrich, T5293) in PBS + 1% BSA. This was followed by three washing steps in PBS + 1% BSA, incubation for 1 hour at 37°C with 1:300 mouse *α*-tubulin antibody (Sigma Aldrich, T6199) diluted in PBS 1% BSA, three more washes in PBS + 1% BSA, incubation for 45 min at 37°C with 1:300 goat anti-mouse AF647 antibody (Life Technologies, A21237) diluted in PBS 1% BSA and three more washes in PBS.

For AF647 heavy chain and light chain clathrin and DNA-PAINT *α*- and *β*-tubulin, the cells were incubated for 1 hour at 37°C with 1:400 mouse anti-light chain clathrin antibody (Sigma Aldrich, C1985) in PBS + 1% BSA and washed three times with PBS + 1% BSA, incubated again for 1 hour at 37°C with 1:400 mouse anti-heavy chain clathrin antibody (Sigma Aldrich, C1860) in PBS + 1% BSA and washed three times with PBS + 1% BSA. Then they were incubated for 45 min at 37°C with 1:400 anti-mouse AF647 antibody (Life Technologies, A21237) in PBS + 1% BSA, washed three times with PBS + 1% BSA, and incubated again for 1 hour at room temperature with 1:400 mouse *β*-tubulin antibody (Sigma Aldrich, T5293) in PBS + 1% BSA. This was followed by three washing steps in PBS + 1% BSA, incubation for 1 hour at 37°C with 1:400 mouse *α*-tubulin antibody (Sigma Aldrich, T6199) diluted in PBS 1% BSA, three more washes in PBS + 1% BSA, incubation for 2 hours at 37°C with 1:100 anti-mouse-D1 Ultivue secondary antibody diluted in antibody dilution buffer (Ultivue) and washed three more washes in PBS.

In any case, after the immunolabeling of tubulin and/or clathrin, a post-fixation step was performed using PBS with 3.6% formaldehyde for 15 min. The cells were washed in PBS three times and then reduced for 10 min with 50 mM NH_4_Cl (Sigma Aldrich, 254134), followed by three additional washes in PBS.

To prepare the neuron samples, rat hippocampal neurons from E18 pups were cultured on 18 mm coverslips at a density of 6,000/cm^2^ according to previously published protocols [29] and following guidelines established by the European Animal Care and Use Committee (86/609/CEE) and approval of the local ethics committee (agreement D18-055-8). After 16 days in culture, neurons were fixed using 4% PFA in PEM (80 mM Pipes, 5 mM EGTA, and 2 mM MgCl_2_, pH 6.8) for 10 min. After rinses in 0.1 M phosphate buffer (PB), neurons were blocked for 60 minutes at room temperature in immunocytochemistry buffer (ICC: 0.22% gelatin, 0.1% Triton X-100 in PB). Following this, neurons were incubated with a chicken primary antibody against map2 (abcam, ab5392) mouse primary antibody against *β*2-spectrin (BD Bioscience, 612563) and a rabbit primary antibody against adducin (abcam, ab51130) diluted in ICC overnight at 4°C, then after ICC rinses with AF 488, 555 and 647 conjugated secondary antibodies for 1h at 23°C.

The *E. coli* K12 (MG1655) cells were grown in 2YT medium (Sigma, Tryptone 16.0 g/L, Yeast extract 10.0 g/L, NaCl 5.0 g/L) at 37°C under agitation (180 rpm). Overnight cultures were diluted 100 times in fresh medium (final volume 300 μL) containing Kdo-N_3_ (1.0 mM). Bacteria were incubated at 37°C for 9 hours under agitation (180 rpm). Then 200 μL of the obtained suspension were washed 3 times with PBS buffer (200 μL, 12,000 rpm, 1 min, room temperature). The pellet was re-suspended in 200 μL of a solution of DBCO-Sulfo-Biotin (JenaBioscience, CLK-A116) (0.50 mM in PBS buffer) and the suspension was vigorously agitated for 90 min at room temperature. Bacteria were washed 3 times with PBS buffer (200 μL, 12,000 rpm, 1 min, room temperature). The pellet was re-suspended in a solution of Streptavidin-AF647 / Streptavidin-AF555 (20 μg/mL each) (Invitrogen, ThermoFischer Scientific, S21374 and S32355) in PBS containing BSA (1.0 mg/mL, 200 μL) and the suspension was agitated at room temperature for 90 min in the dark. Bacteria were then washed 3 times with PBS buffer (200 μL, 12,000 rpm, 1 min, room temperature). The pellet was re-suspended in PBS buffer (400 μL) and stored at 4°C until analysis.

### Fluorescent beads sample preparation

20-nm fluorescent dark red beads samples (**Fig. 2d**, **Supplementary Fig. 5**) were prepared using a 5.10^−7^ dilution of the initial solution (F8783, Thermo Fisher). We performed the dilution in PBS + 5% glucose to match the index of the dSTORM imaging buffer, and we waited for 5 min before starting the acquisition so that the beads had time to deposit on the coverslip.

100-nm diameter tetraspeck fluorescent beads samples (**Supplementary Fig. 4**) were prepared by diluting the initial solution (T7279, Thermo Fisher) at 5.10^−4^ in PBS + 5% glucose, and we waited for 5 min before starting the acquisition for the beads to deposit on the coverslip.

The samples of 40-nm diameter dark red fluorescent beads deposited on a coverslip (**Supplementary Fig. 6a**, **Supplementary Fig. 7**) were obtained by diluting the initial solution (10720, Thermo Fisher) at 5.10^−7^ in PBS + 5% glucose, and we waited for 5 min before starting the acquisition for the beads to deposit on the coverslip.

The samples of 40-nm diameter dark red fluorescent beads randomly distributed in the imaging volume (**Fig. 1c**, **Supplementary Fig. 6b**) were obtained by taking fixed, unlabeled COS-7 cells and adding 500 μL of beads solution (10720, Thermo Fisher) diluted at 5.10^−7^ in PBS during 5 minutes for beads to deposit before removing the solution and replacing it with PBS + 5% glucose. Beads stuck on the upper side of the membrane were thus located at random heights.

### Image acquisition

(d)STORM/DNA-PAINT imaging on biological samples was performed using an oblique epifluorescence illumination configuration. To induce most of the molecules in a dark state, we used either a (d)STORM buffer (Abbelight Smart kit) or a dilution of DNA-PAINT imagers in imaging buffer. In both cases, the sample was lit with an irradiance of approximately 4 kW.cm^−2^ until a sufficient density of molecules was obtained—typically below one molecule per 4 μm^2^ (see **Supplementary Note 1** for a study of the influence of the molecule density per frame on the localization performance). We then started the data acquisition with 50-ms (for AF647) or 100-ms (for AF555 and DNA-PAINT imagers) exposure time and 150 EMCCD gain. The total number of acquired frames was typically between 15,000 and 30,000 per acquisition.

Performance measurements on fluorescent beads were done at low illumination powers (0.15 kW.cm^−2^ for 20-nm diameter dark red beads and 0.025 kW.cm^−2^ for tetraspeck beads and 40-nm diameter dark red beads). The beads were immersed in PBS + 5% glucose and the exposure times and EMCCD gain were 50 ms and 150 respectively. Except for the long-term axial drift tracking experiment, 500 frames were recorded for each performance characterization acquisition.

The acquisition was performed using the Nemo software (Abbelight).

### Localization software

Each 512×512-pixel frame was pre-processed by removing the pixel per pixel temporal median of the previous 10 frames in order to get rid of the slowly varying background without altering the number of photons in the PSFs. The filtered frames were then split in two parts corresponding to the UAF and EPI paths of the DAISY module respectively. On the 512×256-pixel sub-frames, the PSFs were detected using a second order wavelet filtering associated with an intensity threshold (typically 1.0 for the EPI channel, 0.8 for the UAF channel). Each PSF was characterized using a center of mass detection to retrieve the lateral positions *x*^*EPI*^, *y*^*EPI*^, *x*^*UAF*^ and *y*^*UAF*^, and a Gaussian fitting to assess the PSF widths 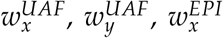 and 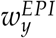. A photon counting was also performed over a 2 μm × 2 μm square area centered on the PSF to determine the number of photons *N*^*EPI*^ and *N*^*UAF*^. A filtering step based on photon numbers (typically 500 photons minimum for AF647), EPI PSF widths 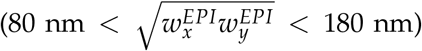 and EPI PSF anisotropy 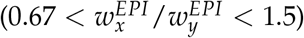 was then operated to get rid of false positive detections. Furthermore, pairs of localizations closer than 2 μm were discarded to avoid biases due to the signal from neighbouring PSFs. Corrections were applied to photon numbers (as mentioned in the **Optical setup** section) and lateral positions *x*^*UAF*^ and *y*^*UAF*^ (to compensate the image deformation induced by the astigmatism as illustrated in **Fig. 2e** and **Supplementary Fig. 7**). Afterwards, the axial positions were calculated: the values of *z*^*SAF*^ were obtained using the theoretical curve provided in [15] whereas those of *z*^*astigmatic*^ could be retrieved by fitting 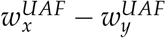 to the calibration curve (see the **Astigmatism calibration** section) using a least squares calculation. Lateral drifts were then corrected using a temporal cross-correlation algorithm. Furthermore, *z*^*astigmatic*^ positions were corrected using the SAF reference (see **Astigmatism correction algorithm** section).

Finally, the values of *z*^*SAF*^ and *z*^*astigmatic*^ were merged together, as well as the values of *x*^*EPI*^ and *x*^*UAF*^, *y*^*EPI*^ and *y*^*UAF*^ (as described in the **Position merging** section).

All this processing was performed using a home-written Python code.

### Astigmatism calibration

Although in the literature, the calibration of axial detection methods is often performed by using fluorescent beads deposited on a coverslip and defocusing the objective, this method is biased since it does not take into account the effect of the spherical aberration, which affects both the position of the focal plane (the so-called focal shift) and the shapes of the PSFs. While the former can be compensated using a calculated correction factor depending on several experimental parameters, there is no simple way to get to correct the latter to our knowledge. Thus we chose to perform the calibration of the astigmatic detection using a sample of known geometry in the nominal acquisition conditions, i.e. with a fixed focus plane and dSTORM fluorophores. More specifically, we used a sample of 15 μm microspheres decorated with fluorophores (either AF647 or AF555), as described in [20]. By measuring the position of the center and the radius of the spheres, it is possible to calculate the expected axial position of each molecule from the measurement of its lateral position. Such an acquisition provides the lookup table giving the correspondence between PSF widths and axial positions.

### Astigmatism correction algorithm

Before combining the two sources of axial information, the astigmatic positions were corrected in order to make them benefit from the SAF absolute detection. This was done thanks to a cross-correlation algorithm between the SAF and astigmatic positions measured for each molecule. As the SAF detection is efficient mostly close to the coverslip, we restricted the data to the subset of molecules verifying *z*^*SAF*^ ∈ [−50 nm, 300 nm] in order to perform the cross-correlation in the domain where both axial information sources are precise and reliable.

First, we removed the tilt: the *z*^*SAF*^ − *z*^*astigmatic*^ axial discrepancy was calculated for each molecule from the data verifying *z*^*SAF*^ ∈ [−50 nm, 300 nm]. The spatially resolved axial discrepancy information was used to calculate the tilt by fitting a plane to the data, which provided the tilt direction and amplitude. The astigmatic positions were corrected in accordance with this result.

Then data was divided in subsets of 1,000 frames and distributed in series of 3D images with 100 nm lateral and 15 nm axial pixel sizes, each of them corresponding to a 1,000 frame subset. For each subset, the SAF and astigmatism 3D images were cross-correlated allowing only axial displacements to maximize the overlap, which brought the correction to be applied to the astigmatic positions for the subset. Then the results obtained for all the subsets were pooled and interpolated to generate the axial drift curve. Thanks to this correction, the astigmatic results were made absolute (i.e. referenced to the coverslip) and insensitive to both the chromatic aberration and the axial drift.

It is worth noting that the 1,000-frame division corresponds to a 50-s sampling of the axial drift (with 50-ms exposure time). This value seems reasonable given the slow evolution of the drift: it is the result of a compromise between the bandwidth of the correction (a finer sampling allows a better correction of higher drift frequencies) and the robustness of the algorithm (if the amount of data is too low, the algorithm may not adequately converge or provide a wrong value). Shorter slices might be used with higher density samples. Similarly, acquisitions featuring a lower SNR or photon number would require larger pixels or larger slices to compensate the influence of the localization precision worsening. The final accuracy of the correction appears to be typically better than 3 nm (this was obtained by measuring the height of fluorophores deposited at the coverslip outside of cells).

### Position merging

In DAISY acquisitions, the lateral positions were obtained by combining the two sources of lateral information according to their uncertainties (the CRLB values were used for that purpose). The exact formula follows the normal distribution combination law:

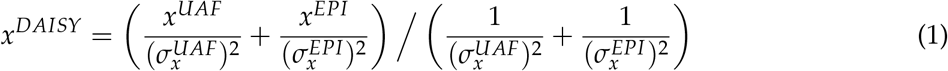

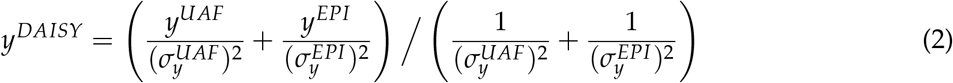

where 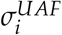 and 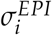 are the localization precisions in the direction *i* for the UAF and EPI detections respectively (i.e. the standard deviations of the positions).

Similarly, the two sources of axial information were merged according to their uncertainties:

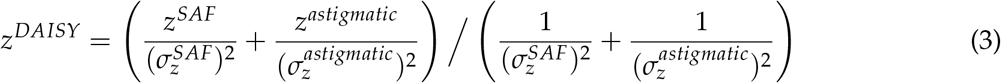

where 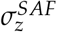 and 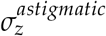 are the axial localization precisions of the SAF and the astigmatic detections respectively.

This combination optimizes the final precision, i.e. it provides the best precision attainable from the two sources given their respective uncertainties.

The relative weights used for DAISY are shown in **Fig. 1b**. It is worth noting that since localization precisions vary with depth, the corresponding weights vary accordingly. Notably, the weight of the SAF detection is more important than that of the axial astigmatic detection at the coverslip, but it quickly dwindles to almost zero after 500 nm. Similarly, the (unastigmatic) EPI detection is more precise in the first depth of field, whereas the (astigmatic) UAF detection dominates after 600 nm, where the EPI PSFs are too defocused to be detected.

### Localization precision measurement

To obtain the localization precisions displayed in **Fig. 1c**, we prepared a sample of 40-nm dark red fluorescent beads randomly distributed in the imaging volume (see **Fluorescent beads sample preparation** section). The results of several 500-frame acquisitions were pooled and for each of them, the lateral drift was corrected. The average axial position was measured for each bead, as well as the standard deviations on the lateral and axial measured positions, which gave the localization precisions. The laser power was adjusted so that the photon numbers emitted by the beads matched those of AF647 (2750 UAF photons per PSF and 2750–5100 EPI photons, depending on the depth of the bead).

Using fluorescent beads seems to be a more reliable method to measure the localization precisions than with biological samples—unlike the fluorescent beads, the use of biological samples require many assumptions on the size and geometry of the labeled target, the label (which is typically around 10–15 nm in the case of immunolabeling), the fluorophore itself, as well as the motion freedom of the label.

### Cramér-Rao Lower Bound calculation

To derive the CRLB for DAISY, we first estimated the lower bounds associated to the astigmatic and the SAF detections separately. To this end, we assumed elliptical Gaussian PSFs for the UAF image and circular Gaussian PSFs for the EPI image. We used a realistic set of parameters corresponding to typical experimental conditions with AF647, i.e. 100 background photons per pixel on each path and a number of photons per PSF equal to 2750 for the UAF path and 2750–5100 for the EPI path (depending on the axial position). The CRLB of the SAF was adapted from [30] and that of the astigmatism was derived from [31]. Finally, the DAISY axial CRLB was obtained from the previous results using **equation** (3). Similarly, the lateral CRLB for the UAF and EPI paths were obtained from [32] and the lateral lower bound of DAISY was calculated from these results using **equations** (1) and (2). See **Supplementary Note 2** for a more exhaustive description of the CRLB calculations. These results were used to plot the curves displayed in **Fig. 1c** and **Supplementary Fig. 1** and **Supplementary Fig. 3**.

Note that the CRLB values are somewhat optimistic and that they are not necessarily expected to be reached in real experimental conditions because they do not account for optical aberrations, polarization effects on the PSF shape or for the ability of the localization algorithm to actually extract the best possible information.

### Data visualization

The 3D view in **Fig. 3b** was obtained using the Nemo software (Abbelight). A filter based on the local density of molecules associated with a threshold was applied on **Fig. 4f–h** to remove false positive detections.

### Code availability

The localization and the lateral drift correction may be performed with any localization software. The DAISY correction code is available on Github at this address: https://github.com/ClementCabriel/DAISYcorrection.

### Data availability

Several localization datasets (data filtered, lateral drift corrected, DAISY axial correction not applied) are available on Github as test samples for the DAISY correction code: https://github.com/ClementCabriel/DAISYcorrection. The authors also uploaded one clathrin-AF647 dataset obtained with DAISY (all corrections performed, data not filtered) on the Shareloc platform: https://shareloc.xyz/#/view?u=z2Dig7bFraDdSHkXwg7Zhv. The authors will keep uploading datasets, both on Github and Shareloc. Other data are available from the corresponding authors upon reasonable request.

## Supplementary material

**Supplementary Figure 1:**
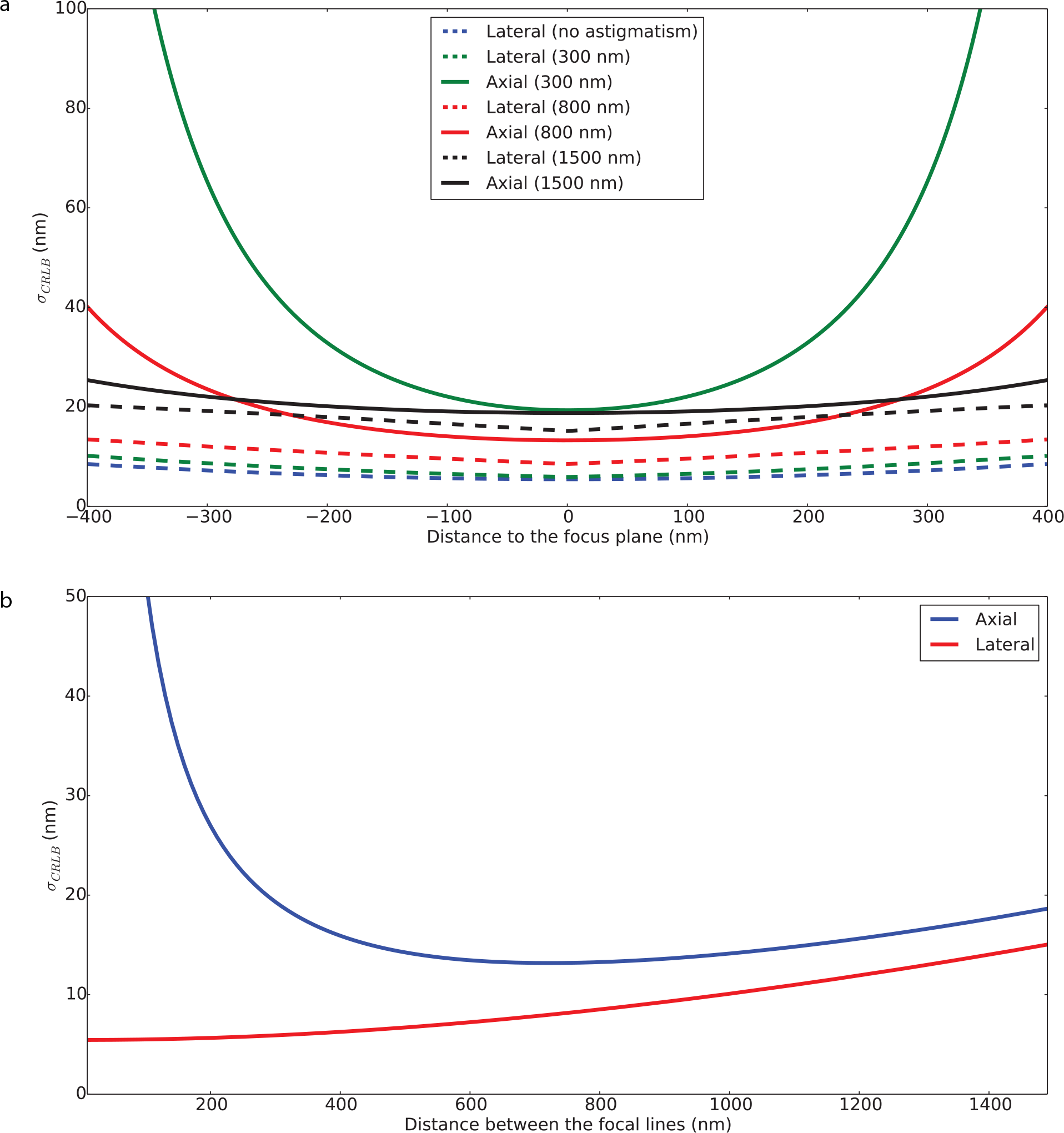
Influence of the depth and the astigmatism amplitude on the axial and lateral Cramér-Rao Lower Bound (CRLB) theoretical limits for 2750 photons per PSF. **(a)** Variation of the localization precision with the distance to the focal plane for different astigmatism amplitudes (expressed as the distance between the two focal lines in the object space, 300 nm being a typical value found in the literature and 800 nm being the value used for DAISY). The solid and dashed lines stand for the axial and lateral precisions respectively. See **Supplementary Note 2** for an explanation of the CRLB calculations. **(b)** Influence of the astigmatism amplitude on the best achievable axial and lateral precisions (i.e. CRLB values at *z* = 0). Note that the axial precision displays a minimum around 800 nm, which we chose as the astigmatism amplitude for DAISY to optimize the axial detection.

**Supplementary Figure 2:**
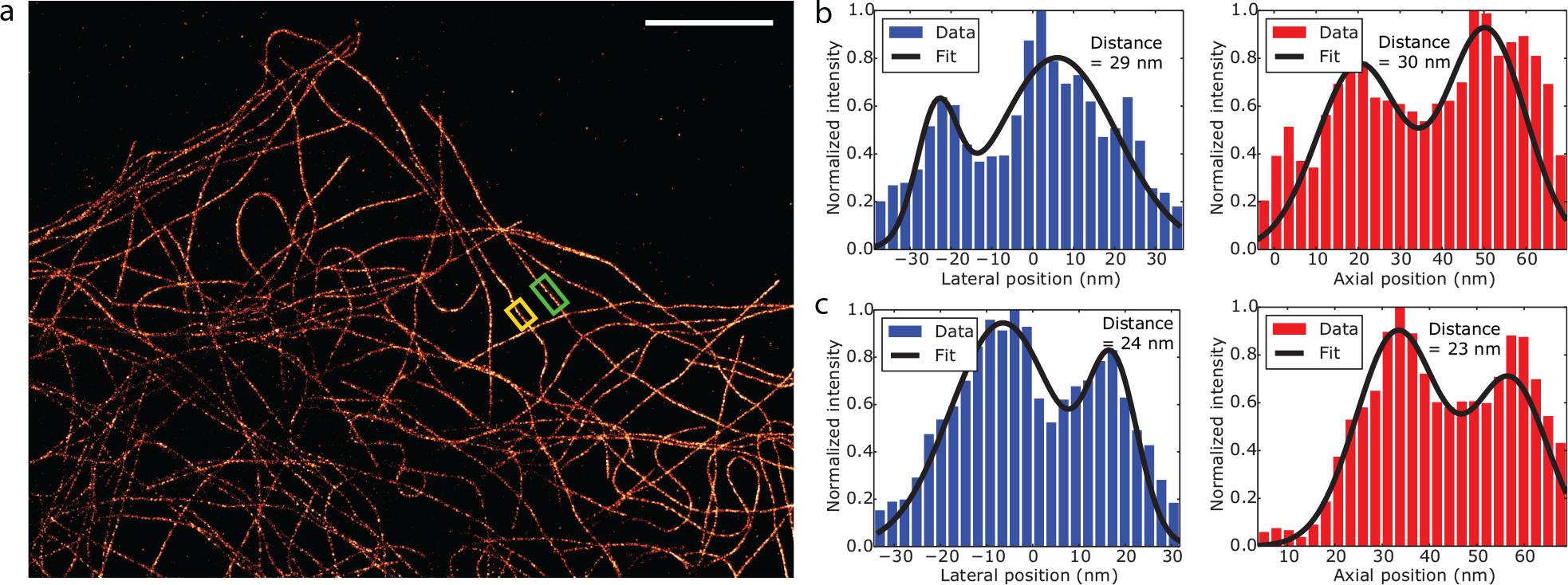
Visualization of the hollowness of microtubules to measure the localization precision. **(a)** 2D super-localized image of COS-7 cells with *α*-tubulin labelled with AF647. The acquisition is the same as that presented in **Fig. 2e**. The lateral (blue) and axial (red) histograms are then plotted in the green boxed region **(b)** and in the yellow boxed region **(c)**. Both the experimental data and the fitted profile with a double Gaussian function are displayed, as well as the distance between the two Gaussian peaks. The hollowness is clearly visible, and the distance between the peaks corresponds to a localization precision around 14–16 nm (see Ref. [1], **Supplementary Fig. 7**). Scale bar: 5 μm.

**Supplementary Figure 3:**
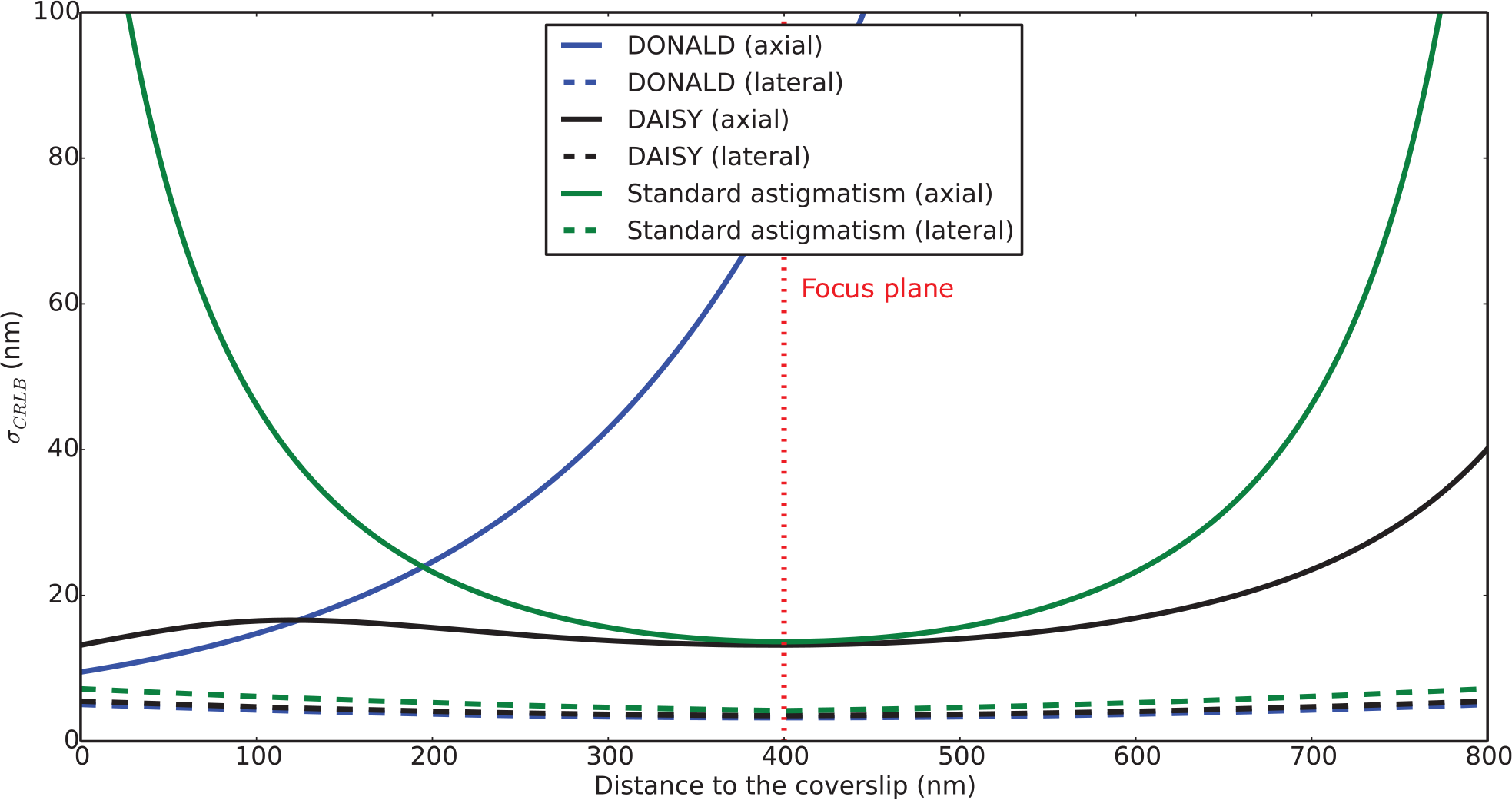
Comparison of the lateral and axial CRLB for DONALD, standard astigmatism and DAISY as a function of the depth. The number of photons is 2750 for the UAF PSFs and 2750–5100 (depending on the axial position) for the EPI PSFs (similar to AF647), and the standard astigmatism corresponds to a 300-nm spacing between the two focal lines. The focus position is assumed to be 400 nm above the coverslip (typical experimental value), which is represented by the red dotted line. The solid and dashed lines stand for the axial and lateral precisions respectively. See **Supplementary Note 2** for an explanation of the CRLB calculations.

**Supplementary Figure 4:**
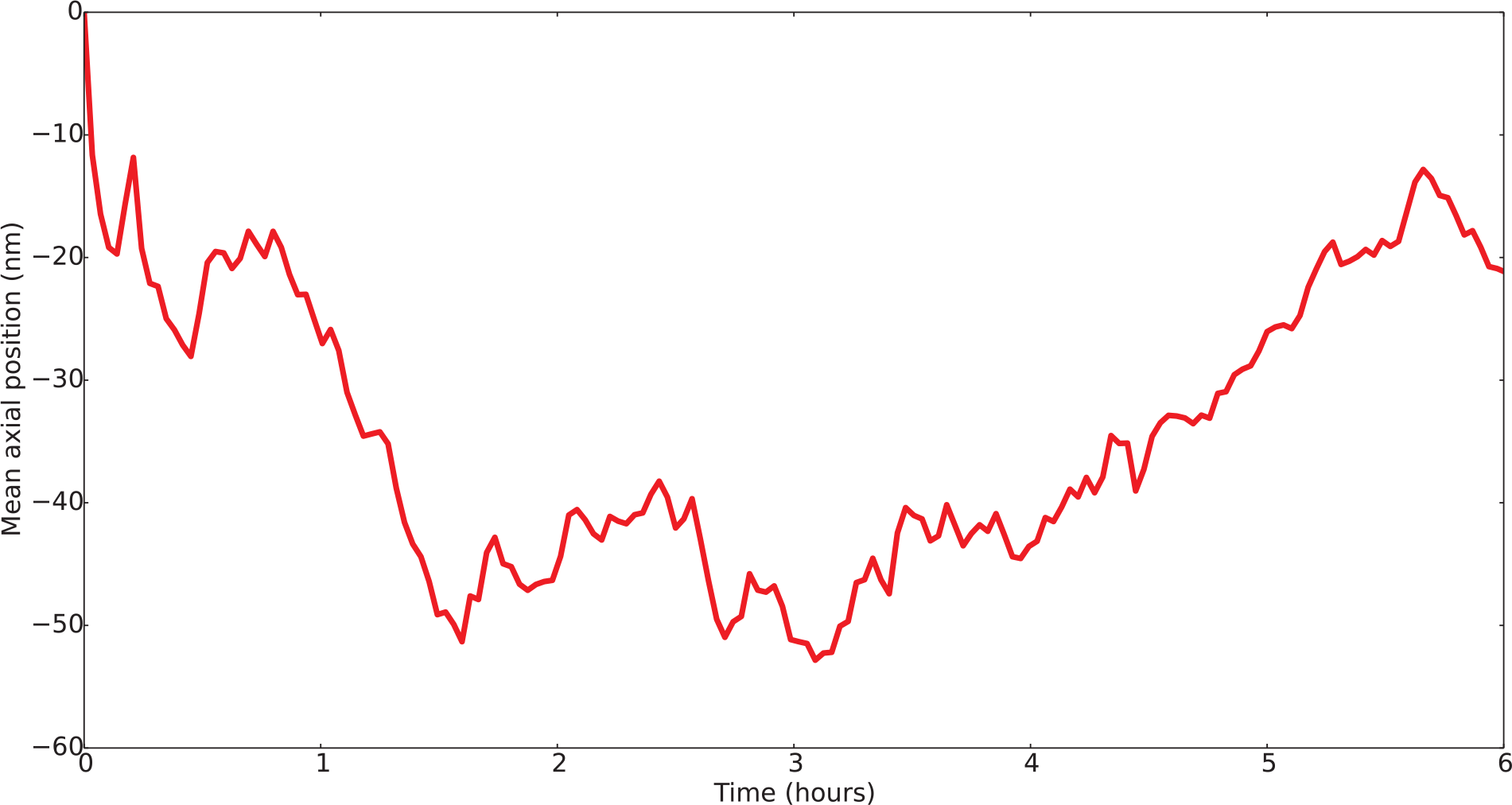
Long-term tracking of the axial drift. The mean axial position of 100 nm diameter tetraspeck fluorescent beads (Thermo Fisher, T7279) over the imaged field is plotted as a function of time over approximately six hours. The results were averaged over 50 frames (i.e. 2.5 seconds) to suppress the influence of the localization uncertainty.

**Supplementary Figure 5:**
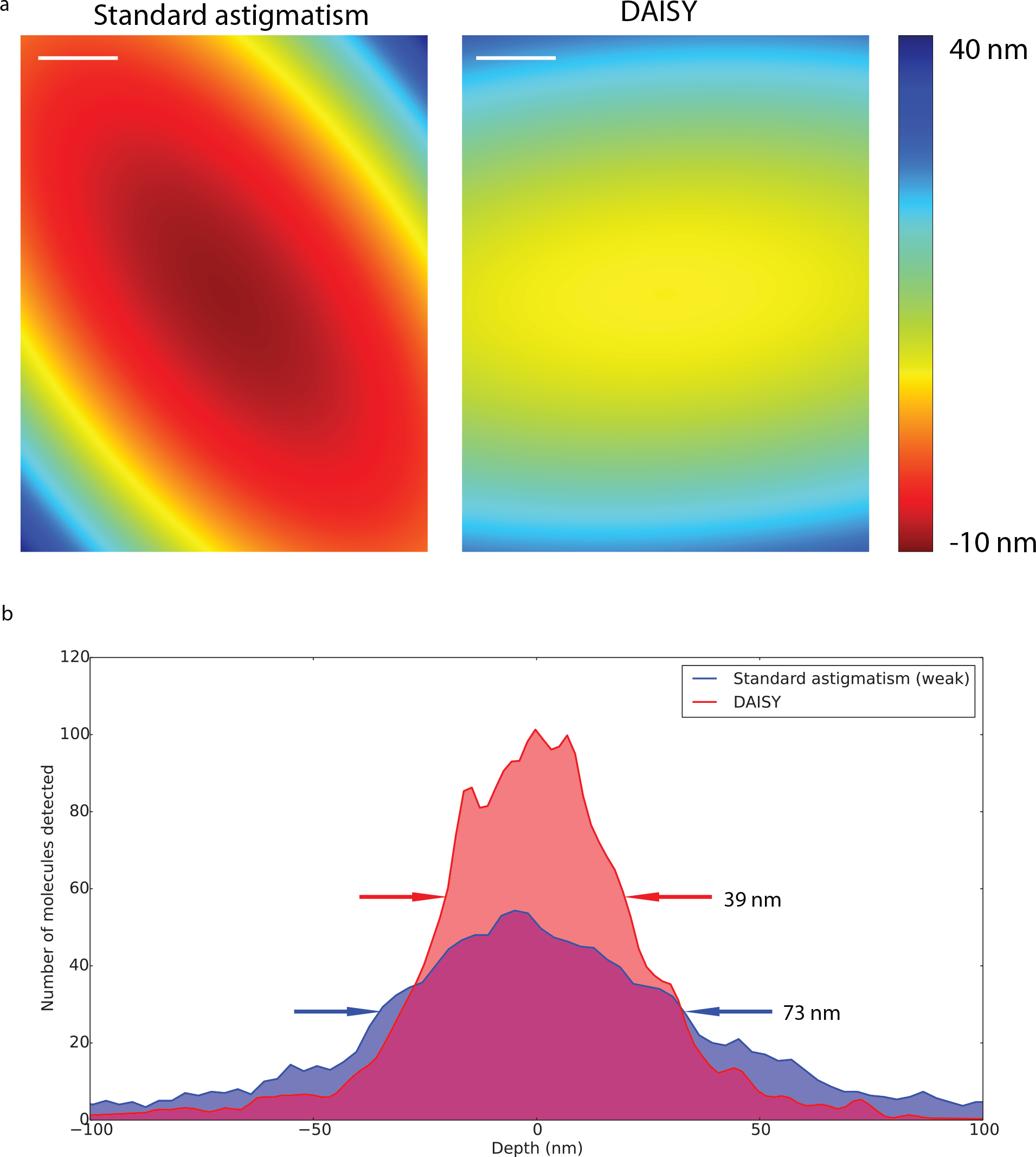
Influence of the remaining field aberrations on the axial detection after tilt correction. **(a)** Interpolated depth maps of the axial positions measured for a sample of 20 nm dark red fluorescent beads deposited on a coverslip and averaged over 500 frames to suppress the influence of the localization precision. The results are plotted for both a typical astigmatism-based imaging (300 nm spacing between the two focal lines, close to the values encountered in the literature) and for DAISY. **(b)** The depth histograms are plotted over the 25-μm wide field for both the typical astigmatic detection (300 nm between the two focal lines) and for DAISY. The displayed widths stand for the full widths at half maximum. Scale bars: 5 μm.

**Supplementary Figure 6:**
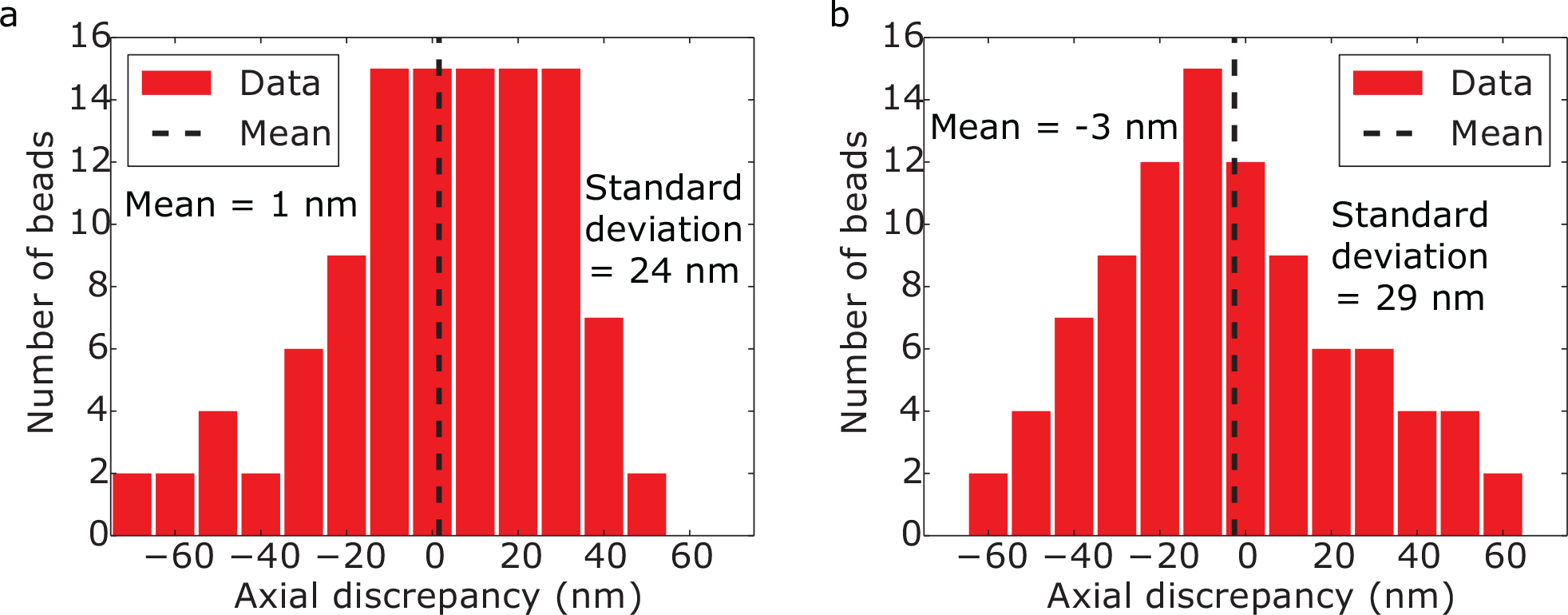
Measurement of the residual axial registration error after the correction. **(a)** Histogram of the residual axial discrepancy (i.e. *z*^*astigmatic*^ − *z*^*SAF*^) obtained with 40-nm diameter dark red fluorescent beads deposited at the coverslip. **(b)** Histogram of the residual axial discrepancy (i.e. *z*^*astigmatic*^ − *z*^*SAF*^) obtained with 40-nm diameter dark red fluorescent beads randomly distributed in the volume and imaged over a 500-nm range above the coverslip capture range. In both cases, the axial positions were averaged over 500 frames to mitigate the influence of the localization precision.

**Supplementary Figure 7:**
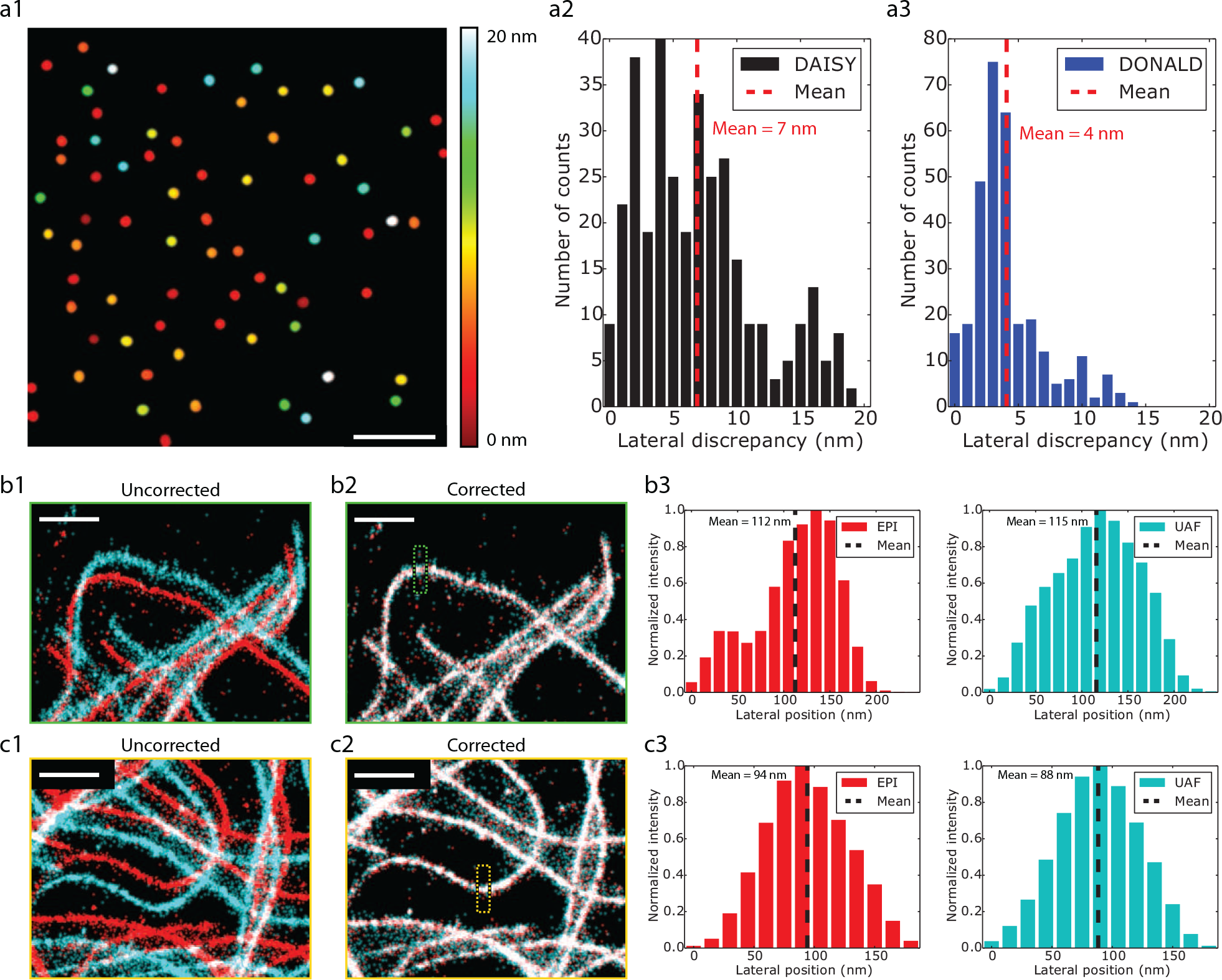
Measurement of the residual lateral registration error after the correction. **(a)** Measurement on 40-nm diameter dark red fluorescent beads deposited at the coverslip (three acquisitions on different fields were stacked): **(a1)** Map of the residual lateral error, **(a2)** Residual lateral error histogram obtained with DAISY. As a comparison, the same measurement is provided for a DONALD acquisition (i.e. the same dual-view setup without the cylindrical lens) in **(a3)**. In both cases, the axial positions were averaged over 500 frames to mitigate the influence of the localization precision. The residual discrepancies are slightly superior for DAISY (7 nm) than for DONALD (4 nm). We attribute this difference to either a PSF shape-dependent lateral bias (which would be larger for aberrated PSFs), or a residual influence of the localization uncertainty. **(b–c)** Measurements performed on the two microtubules regions of interest presented in **Fig. 2e**: **(b1–c1)** Superimposed 2D maps (red: EPI path, cyan: UAF path) before running the correction algorithm. The images display large discrepancies (around 500 nm) due to the magnification difference between the *x* and *y* axes. **(b2–c2)** Superimposed 2D maps after running the correction, **(b3–c3)** EPI and UAF profiles along the axes displayed in **(b2–c2)** after correction. The residual discrepancy is below 6 nm. Scale bars: 5 μm **(a)**, 1 μm **(b)**.

**Supplementary Figure 8:**
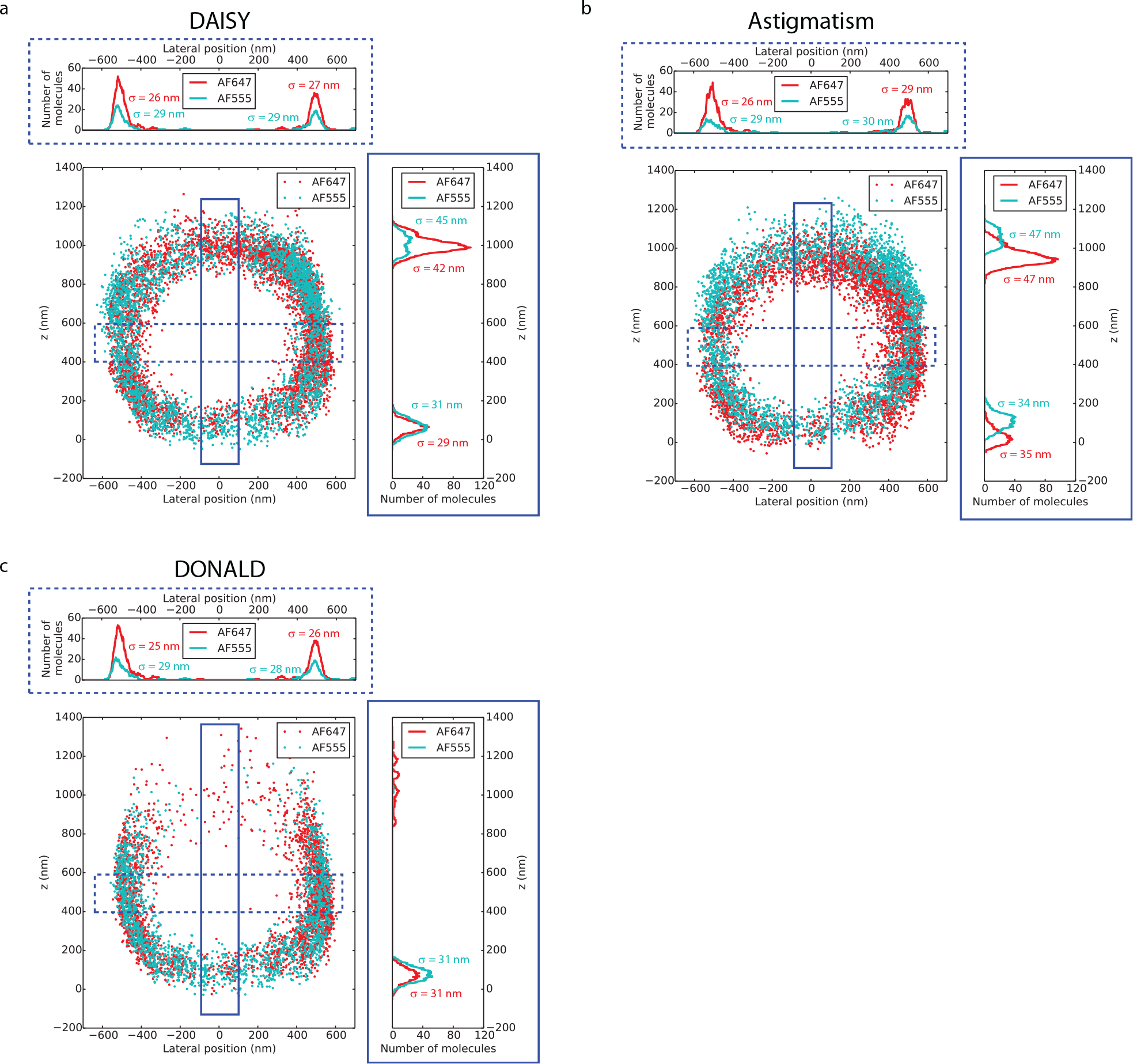
Comparison of the 3D performances of **(a)** DAISY, **(b)** uncorrected astigmatism and **(c)** DONALD on a sample of living *E. coli* bacteria labeled with AF647 and AF555 at the membrane (see **Fig. 3a–c** and **Methods**). The *x*-*z* slices along the line displayed in **Fig. 3a** and the axial and lateral profiles in the boxed regions are plotted. The *σ* values stand for the standard deviation of the distributions. Like DAISY, DONALD features an absolute detection, unsensitive to both chromatic aberration and axial drift. However, the axial precision deteriorates sharply with the depth due to the decay of the SAF signal; thus the top half of the sample (beyond 500 nm) is hardly visible. Uncorrected astigmatism has the same capture range as DAISY, but since it lacks the absolute information, it exhibits an axial shift between the two colors as well as a broadening of the histogram widths due to the axial drift.

**Supplementary Figure 9:**
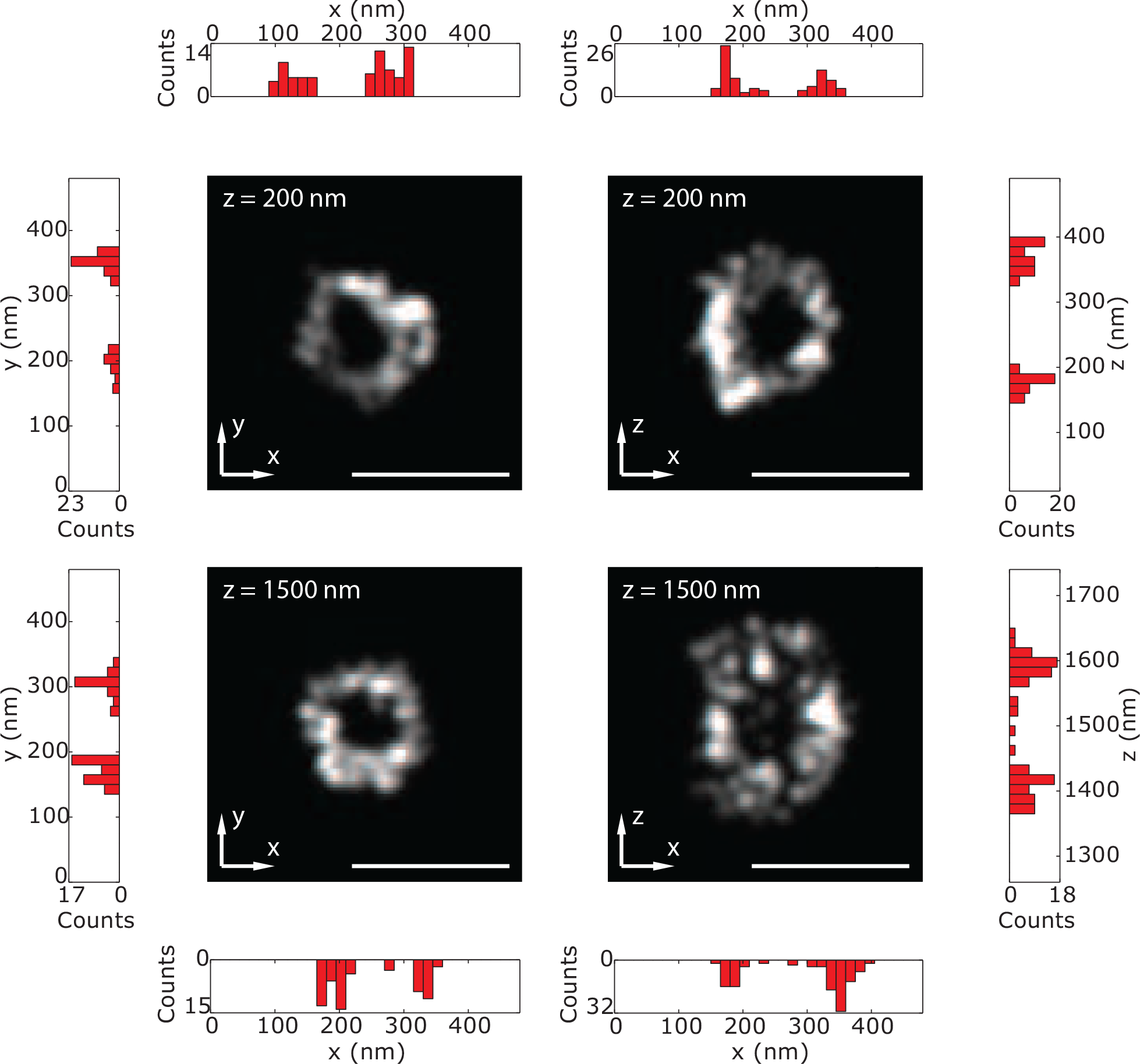
Lateral and axial histograms plotted on the clathrin spheres presented in **Fig. 4g–h**. The histograms are plotted along lines at the center of the spheres to highlight their hollowness. Scale bars: 250 nm.

### Supplementary Note 1: Influence of the molecule density per frame on the localization computation

To assess the impact of the molecule density per frame on the localization performance of our algorithm and filtering, we simulated DAISY PSFs on a 30 μm × 30 μm field with 100 nm pixels. The PSFs were simulated as elliptical gaussians for the UAF channel and circular gaussians for the EPI channel with realistic PSF sizes matching those of experimental data obtained with AF647. All the sources were considered at the coverslip (*z* = 0 nm) and all the PSFs had the same intensity (2750 UAF photons, 2750–5100 EPI photons depending on the axial position of the source). In order to decouple the effect of the molecule density from the localization precision, we did not add background or Poisson noise. The lateral positions were uniformly distributed over the field. The number of PSFs per frame ranged from 1 (1.1 10^−3^ molecules/μm^2^) to 1000 (1.1 molecules/μm^2^). The generated positions were recorded for later use.

Then the localization was run on the generated data and the filters (size and anisotropy of the unastigmatic EPI PSFs, distance between PSF neighboring pairs) were applied and the number of remaining detections was compared to the number of generated molecules to calculate the fraction of missed/discarded localizations (**Supplementary Fig. 10a**). Among the remaining localizations, those displaying a 3D distance to the expected position superior to 50 nm were flagged as wrong detections and their fraction among all the localizations after filtering was displayed in **Supplementary Fig. 10a** in the cases of the SAF and astigmatic detections. Finally, the lateral and axial (for the SAF and the astigmatic detections) median distances to the expected positions were displayed in **Supplementary Fig. 10b**.

As expected, the numbers of missed/discarded localizations and wrong detections increase with the density, but the latter remains quite low for reasonable densities (under 15 % below 0.3 molecules/μm^2^). Similarly, the lateral and axial position discrepancies increase with the density. Realistic dSTORM conditions correspond to densities around 10^−2^–10^−1^ molecules/μm^2^. At such densities, the number of missed/rejected localizations can account for up to 40 % of the total number of localizations, but the number of wrong detections remains minimal (below 4 %). Besides, the errors on the measured positions are rather low (inferior to 4 nm in the lateral and 1 nm in the axial direction, both in SAF and astigmatism).

These results can prove useful to optimize the acquisition conditions—especially the composition of the imaging buffer in dSTORM, the concentration of imager strands in DNA-PAINT or the activation power in PALM, as well as the exposure time for all these methods. In DNA-PAINT acquisitions, relatively high molecule densities (up to 0.3 molecules/μm^2^) can be used to speed up acquisitions, as long as the localization error remains below the localization precision. On the contrary, in PALM experiments, the number of photoactivable molecules is often low and in order to minimize the number of missed/discarded molecules, the molecule density should be kept low (inferior to 3 10^−2^ molecules/μm^2^). Depending on the sensitivity of the fluorophores used to photobleaching, dSTORM acquisitions can match either of the two previously described cases.

**Supplementary Figure 10:**
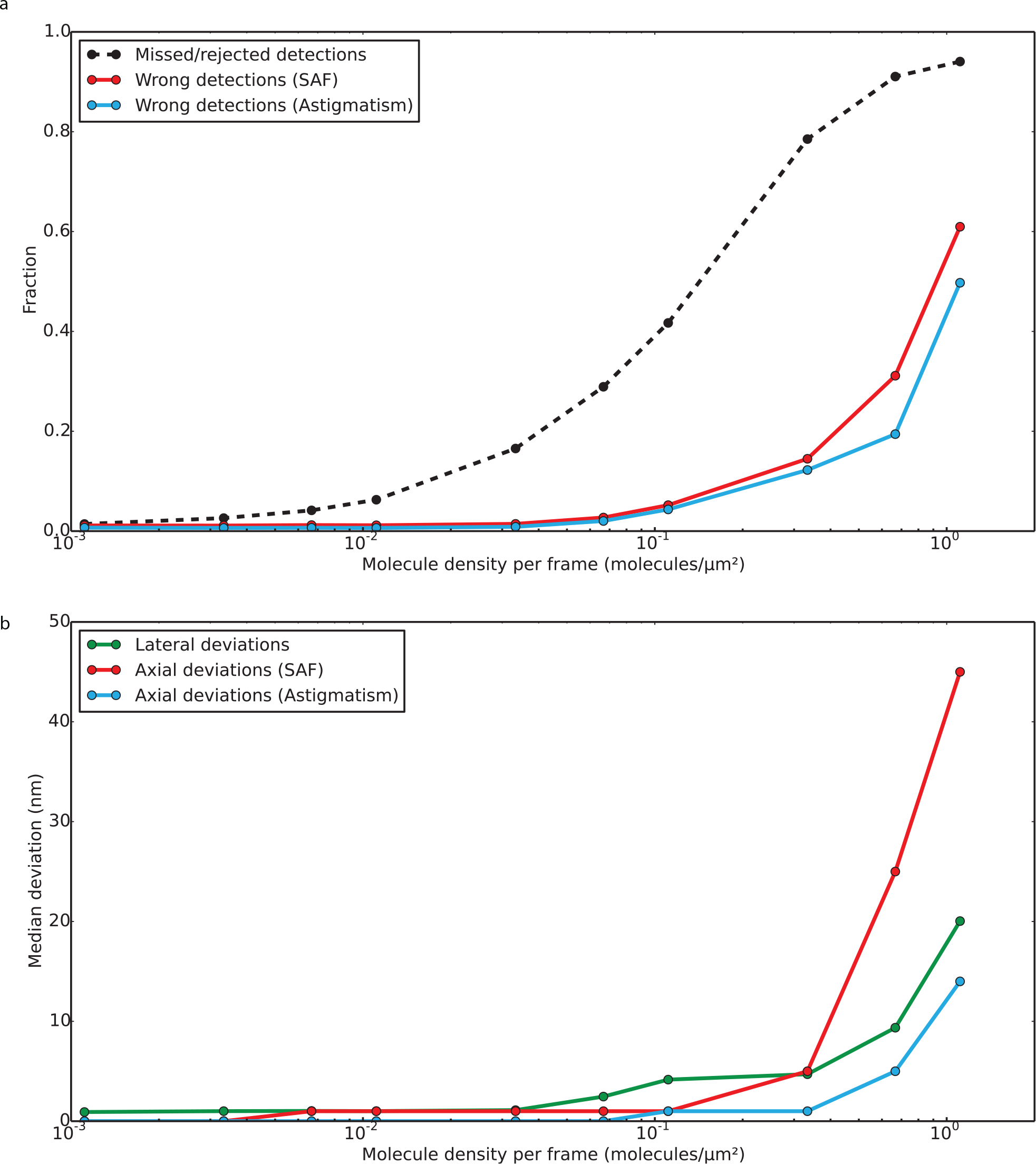
Influence of the molecule density per frame on the localization computation. **(a)** Fraction of localizations missed or discarded (black dashed line) and fraction of wrong detections (blue and red solid lines) as a function of the molecule density on each frame. **(b)** Median lateral and axial discrepancies between the real and the measured positions as a function of the molecule density on each frame.

### Supplementary Note 2: Fisher information and Cramér-Rao Lower Bounds

To determine the theoretical limits of our method, we calculated the Fisher information and the Cramér-Rao Lower Bounds (CRLB) of both the SAF and astigmatic axial detections, as well as that of the lateral detection in order to have access to the theoretical limits of DAISY.

#### 1 Fisher information and CRLB for SAF

We used the same approach as Balzarotti *et al.*[2] to calculate the Fisher information for the supercritical angle fluorescence signal and the associated Cramér-Rao Lower Bounds. Considering an emitter at the position 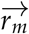 exposed to *K* different illuminations, each photon acquired *n*_*i*_ (*i* ∈ [0, *K* − 1]) follows a Poissonian statistics with a mean *λ*_*i*_ that depends on the illumination. The authors demonstrated that the components of the parameter vector with negligible dark count can be expressed as:

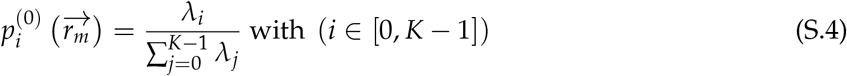

Adding the background signal, this becomes:

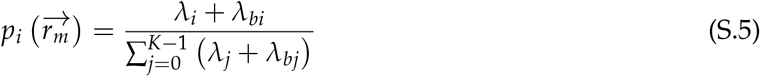

This equation can be simplified:

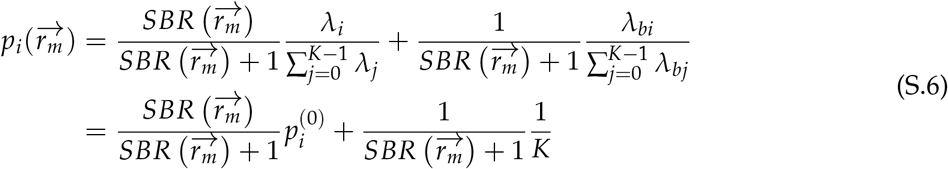

where 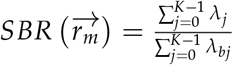 represents the signal to background ratio. Balzarotti *et al.* showed that the Fisher matrix can be expressed in a simple form:

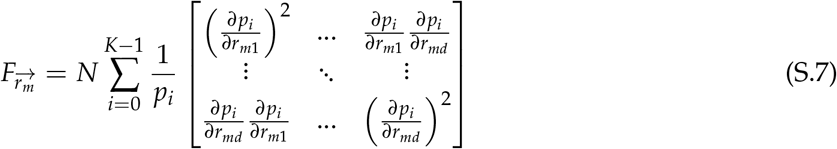

The Fisher information matrix gives access to a lower bound for the covariance matrix 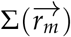. The arithmetic mean of the eigenvalues 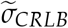 of the lower bound matrix is interpreted as a performance metric:

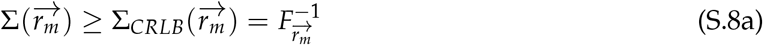

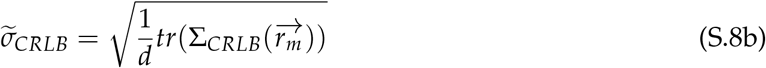

where *d* is the number of dimensions considered and *N* is the total acquired photon number.

These results can be transposed to our model provided a few modifications. Rather than considering different illuminations, we consider a sampling of the signal in two parts: one (EPI signal, noted *i* = 0) dependent on the *z* position of the emitter and one (UAF signal, noted *i* = 1) independent of *z*. In this case, the Fisher matrix takes the form of a scalar:

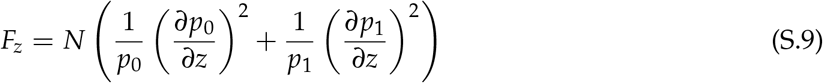

*p*_*i*_(*z*) is provided by (S.6):

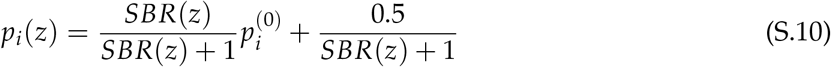

(S.6) can be differentiated:

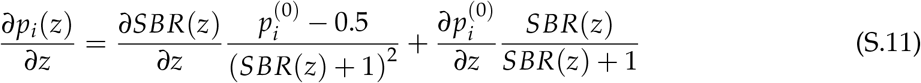

First, we use the theoretical dependence of the SAF signal versus the *z* position by performing simulations based on the work of Wai Teng Tang *et al.* [3]. By fitting the simulation results, we assume that the ratio between the SAF and UAF photon numbers can be approximated as follows for a numerical aperture of 1.49 and an fluorescence wavelength *λ*_*fluo*_:

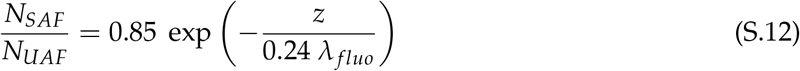

The signal of an emitter is divided in two parts so as to separate the UAF from the EPI fluorescence. In this case, the mean of the Poisson distribution for each part can be expressed as:

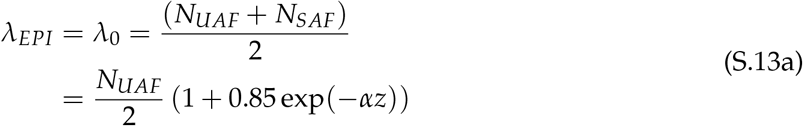

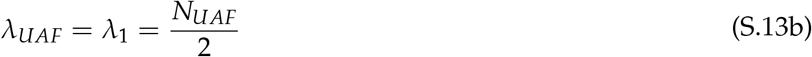

with 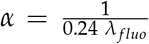. Thses terms can be used in **equation** (S.4) to obtain the two components of the parameter vector with neglected Gaussian noise:

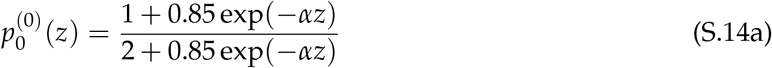

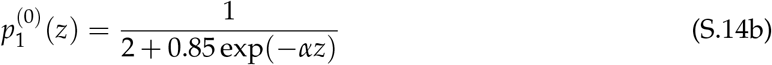

At this point, a background noise term *B* has to be introduced in the calculation. *B* is a photon number associated to an optical signal produced mainly by fluorescent probes located outside the focal plane and is approximated to 200 photons per channel in our calculations. *B* represents *λ*_*bj*_ = *λ*_*b*_, considered constant for each channel. We define the *SBR*(*z*) as :

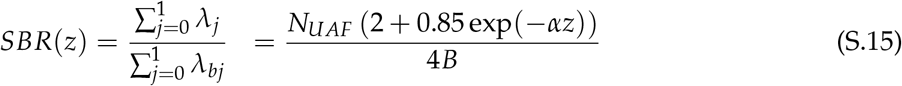

Finally, we can extract the expression of the Fisher information and the CRLB:

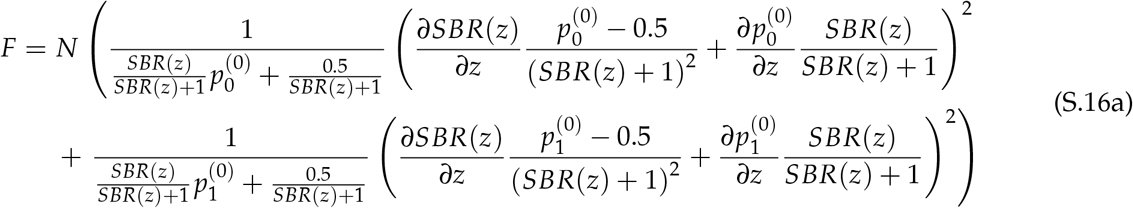

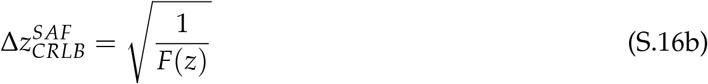

with the different parameters:

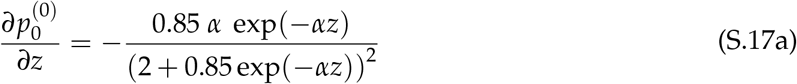

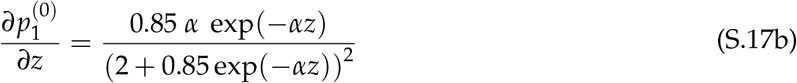

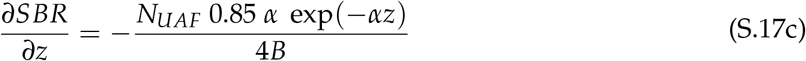

#### 2 CRLB for astigmatism

The Cramér-Rao Lower Bound for the astigmatic detection is directly computed from the work of Rieger and Stallinga [4]. We consider that an astigmatic PSF can be approximated by an elliptical Gaussian PSF with different widths in *x* and *y*, noted *w*_*x*_ and *w*_*y*_:

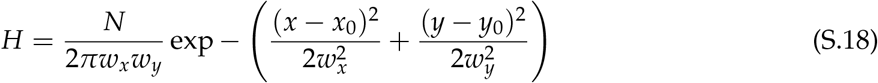

From (S.18), the CRLB for the *w*_*x*,*y*_ parameters can be approximated with the semi-exact formula:

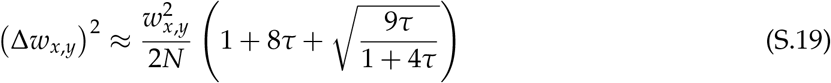

where *τ* is approximately equal to the ratio between the peak and background intensities (*a* being pixel size):

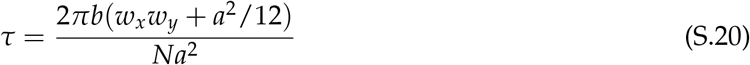

The authors derive the axial detection position from the focus S curve:

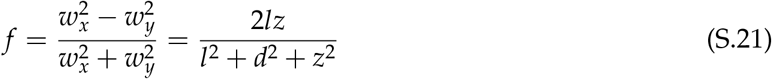

where *d* stands for the focal depth and 2*l* is the distance between the focal lines. Usually, these two parameters are obtained by experimental measurements. The CRLB for the axial position is expressed as follows:

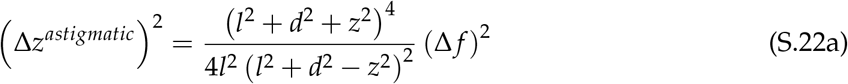

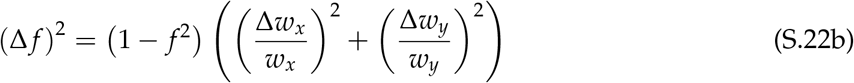

By combining (S.19), (S.21), (S.22a) and (S.22b), the final expression of the CRLB for the axial position of astigmatic method reads:

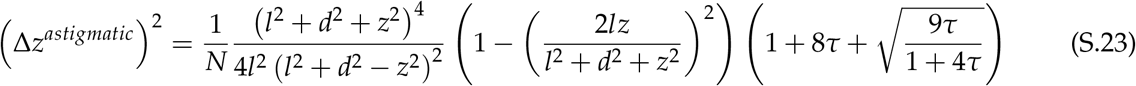

#### 3 CRLB for DAISY

In DAISY, the axial positions from SAF and astigmatism are merged according their uncertainties in order to optimize the final precision (see **Methods**, **Position merging** section, **equation** (3)):

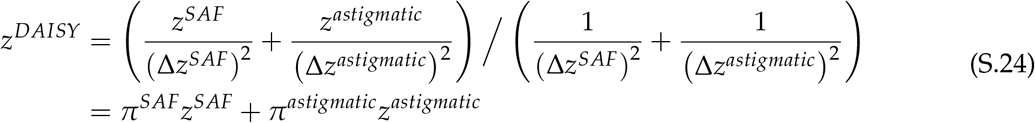

where *π*^*SAF*^ and *π*^*astigmatic*^ are the relative weights of the SAF and astigmatic information sources (note that these weights vary with the axial position):

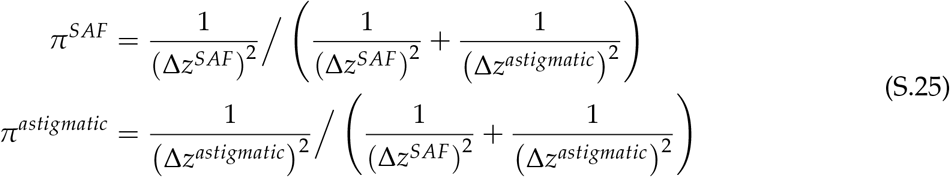

The CRLB for DAISY then reads:

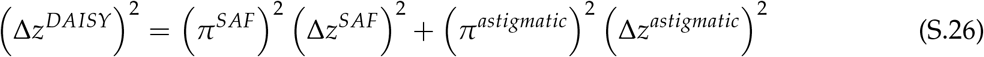

#### 4 CRLB for the lateral detection

The lateral lower bound was obtained using the same assumptions (PSF shape, photon number, background) than those described for the axial detection. We used the formula provided in [5]:

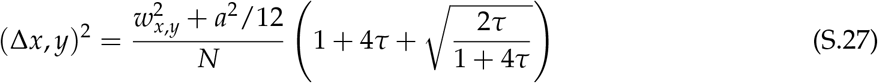

with *τ* defined as in **equation** (S.20).

Like the axial position, the lateral position results from the merging of the measured lateral UAF and EPI positions (see **Methods**, **Position merging** section, **equations** (1) and (2)). Thus it can be written as a weighted sum:

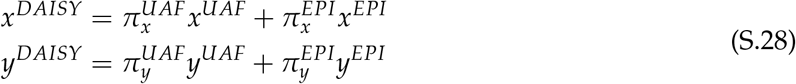

where 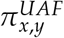 and 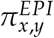 are the relative weights of the UAF and EPI information sources for the *x* and *y* positions respectively (note that these weights vary with the axial position):

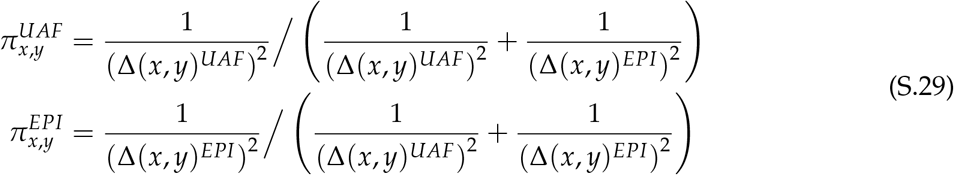

As a result, the CRLB finally reads:

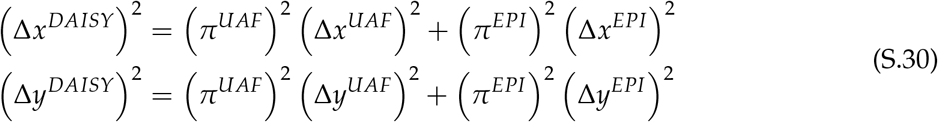

